# Dopamine drives a positive reward bias on human reinforcement learning

**DOI:** 10.64898/2025.12.24.696333

**Authors:** Arnaud Zalta, Vasilisa Skvortsova, Samuel R. Hewitt, Michael Moutoussis, Matthew M. Nour, Raymond J. Dolan, Charles Findling, Tobias U. Hauser, Valentin Wyart

## Abstract

Formal theories of reinforcement learning (RL) prescribe a clearly defined function for dopamine, namely modulating learning via reward prediction errors (RPEs). Yet, empirical evidence in humans remains scarce, and recent advances introducing noisy RL cast doubt on a simple one-to-one mapping between neurotransmitters and computational mechanisms. Here, we detail a double-blind, placebo-controlled, randomised pharmacological study using the dopamine precursor L-DOPA, while healthy volunteers performed a volatile two-armed bandit task. Behaviourally, L-DOPA decreased switching behaviour following below-average rewards. Algorithmic RL modelling of human behaviour supported a dual effect of L-DOPA on the rate and precision of learning. By leveraging recurrent neural networks (RNNs) as implementational models of RL, we explain this dual effect through a single inference-time modulation, whereby L-DOPA triggers a positive reward bias at the input of the recurrent layer that implements RL. Our findings highlight a unifying mechanism at the implementation level that explain seemingly disparate algorithmic effects of dopamine.

## Introduction

Learning the value of actions helps us adapt in changing or uncertain environments, such as navigating a metro at rush hour or finding our way in an unfamiliar city^1^. Algorithmic computational temporal difference-based formulations of reinforcement learning (TD-RL) provide a powerful account of human learning in such contexts^2,3^. Central to these models is the reward prediction error (RPE) – the difference between the obtained reward and the expected value of the chosen action – that drives action value updates and guides the selection of new actions according to a specific policy. In partially observable environments, previous TD-RL research has explained human behavioural variability as a policy trade-off between exploiting known options and exploring novel one^3–5^. However, recent work shows that a limited computational precision during TD-RL value updates (i.e., noisy learning) acts as a significant contributor to human behavioural variability^6^.

Dopamine is critical for reward-guided learning^7–9^, and seminal studies have shown that transient midbrain dopaminergic activity encodes RPEs^10,11^. But dopamine has also been linked to computations occurring downstream of TD-RL^12^. Indeed, dopaminergic interventions can alter specific forms of exploration^13,14^, and computational models suggest a role of dopamine in shaping action policies^15,16^. Moreover, recent studies show that while transient dopamine release is linked to RPE computation, it is independent of learning rate^17^ – the TD-RL parameter governing how strongly RPEs trigger action value updates. Therefore, the precise computational role of dopamine during reward-guided learning remains unclear, all the more so in humans where the study of neural circuit mechanisms underlying computations in reward-guided learning is particularly challenging. Formal implementational models, such as recurrent neural networks (RNNs), offer a promising approach to link dopaminergic interventions with precise algorithmic TD-RL computations in healthy participants.

In this study, we administered sub-therapeutic doses of dopaminergic medication L-DOPA to healthy adult volunteers in a placebo-controlled, double-blind, randomized study while they performed a restless two-armed bandit task with partially observable outcomes. Behaviourally, L-DOPA decreased switching between options following below-average rewards. Fitting a noisy TD-RL model to participants’ behaviour^6^, we found that L-DOPA decreased the rate and precision of human TD-RL without affecting their action policy. We trained noisy^6,18^ RNNs^19,20^ on the same task as participants and fitted inference-time modulation parameters to the observed effects of L-DOPA on human behaviour. A positive reward bias at the input of the recurrent layer explained the dual influence of L-DOPA on human reward-guided learning. By combining pharmacological protocols in healthy humans with algorithmic and implementational models of RL, we demonstrate that the dual effect of dopamine on human TD-RL can be explained by an upstream bias on the perception of obtained rewards.

## Results

### Dopamine increases tendency to pursue sub-average rewards

Fifty-eight healthy adult volunteers performed a two-armed bandit task after receiving a single oral dose of either Madopar CR (125 mg of L-DOPA + 25mg of Benserazide – Controlled-Release) or L-ascorbic acid (500mg placebo) in the context of a double-blind randomised, between-subjects study (**Fig. 1a**). During the experiment, the participants were instructed to maximize their monetary payoff by repeatedly sampling from one of two bandits, represented by coloured shapes (**Fig. 1b**, see **Methods**). The payoffs from each shape were drawn from probability distributions with means that drifted independently across trials, requiring participants to track these changing mean values over time (**Fig. 1c**).

**Figure 1.**
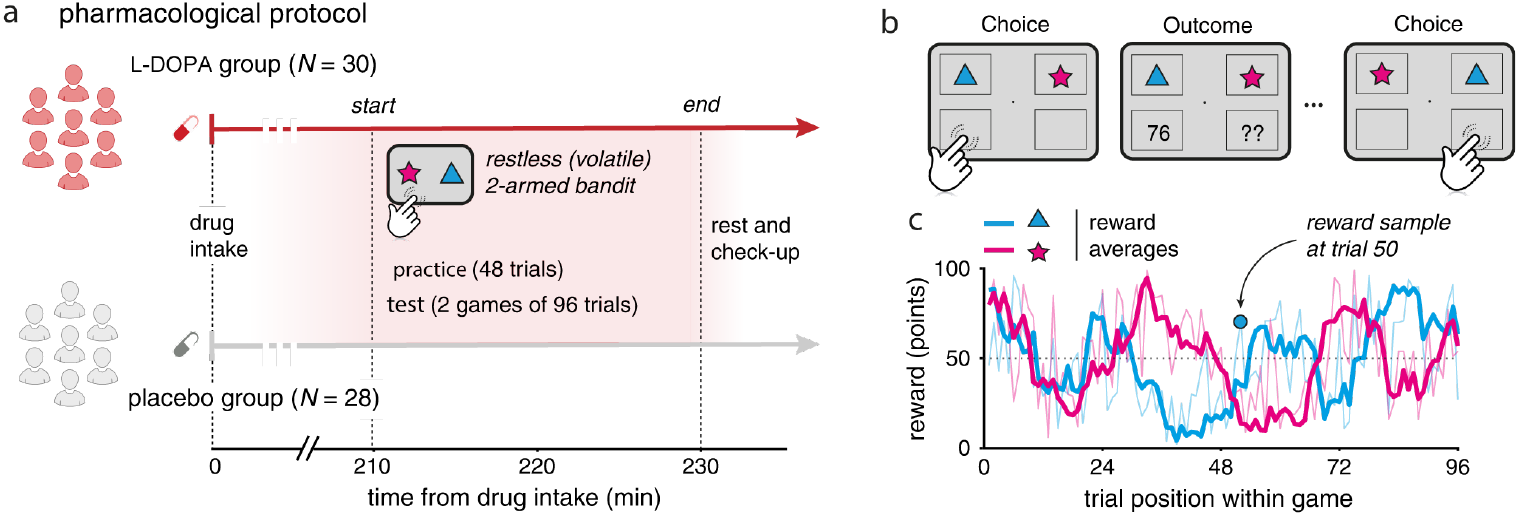
Healthy participants performed a volatile two-armed bandit task under L-DOPA. **a**. Description of the pharmacological protocol. Participants performed a two-armed bandit task and were randomly allocated to one of two drug conditions. Thirty participants performed the task under L-DOPA provided orally, the other twentyeight participants performed the same task under placebo. **b**. Trial structure of the volatile two-armed bandit task. In each trial, participants were asked to choose one of two reward sources depicted by coloured shapes and then observed its associated outcome (from 1 to 99 points, converted into a financial bonus at the end of the experiment). **c**. Example of drifts in the magnitude of rewards that can be obtained from the two sources. Rewards were sampled from probability distributions with means that drifted independently across trials. Thick lines represent the drifting means of the two probability distributions, whereas thin lines correspond to reward samples drawn from the probability distributions that can be obtained if chosen in each trial.

We first assessed whether the drug affected overall performance. We did not observe any effect of L-DOPA on participants’ frequency of choosing the currently best option (i.e., the option with current highest reward mean; rank-sum test: *z* = 1.49, *p* = 0.14; **Fig. 2a**). As expected, we found across both groups that the probability of choosing the same arm as in the previous trial increased as a function of the reward obtained at the previous trial (mixed-effects ANOVA, F (7,399) = 215.2, *p* < 0.001; **Fig. 2b**). This indicates that subjects meaningfully took previous outcomes into account and adjusted their choice behaviour accordingly. Interestingly, the placebo and L-DOPA groups showed differences in this ‘repetition’ curve (mixed-effects ANOVA, main effect of drug: F (1,57) = 16.0, *p* <0.001), an effect that depend on the magnitude of the obtained reward (interaction between drug and obtained reward: F (7,399) = 3.02, *p* < 0.01).

**Figure 2.**
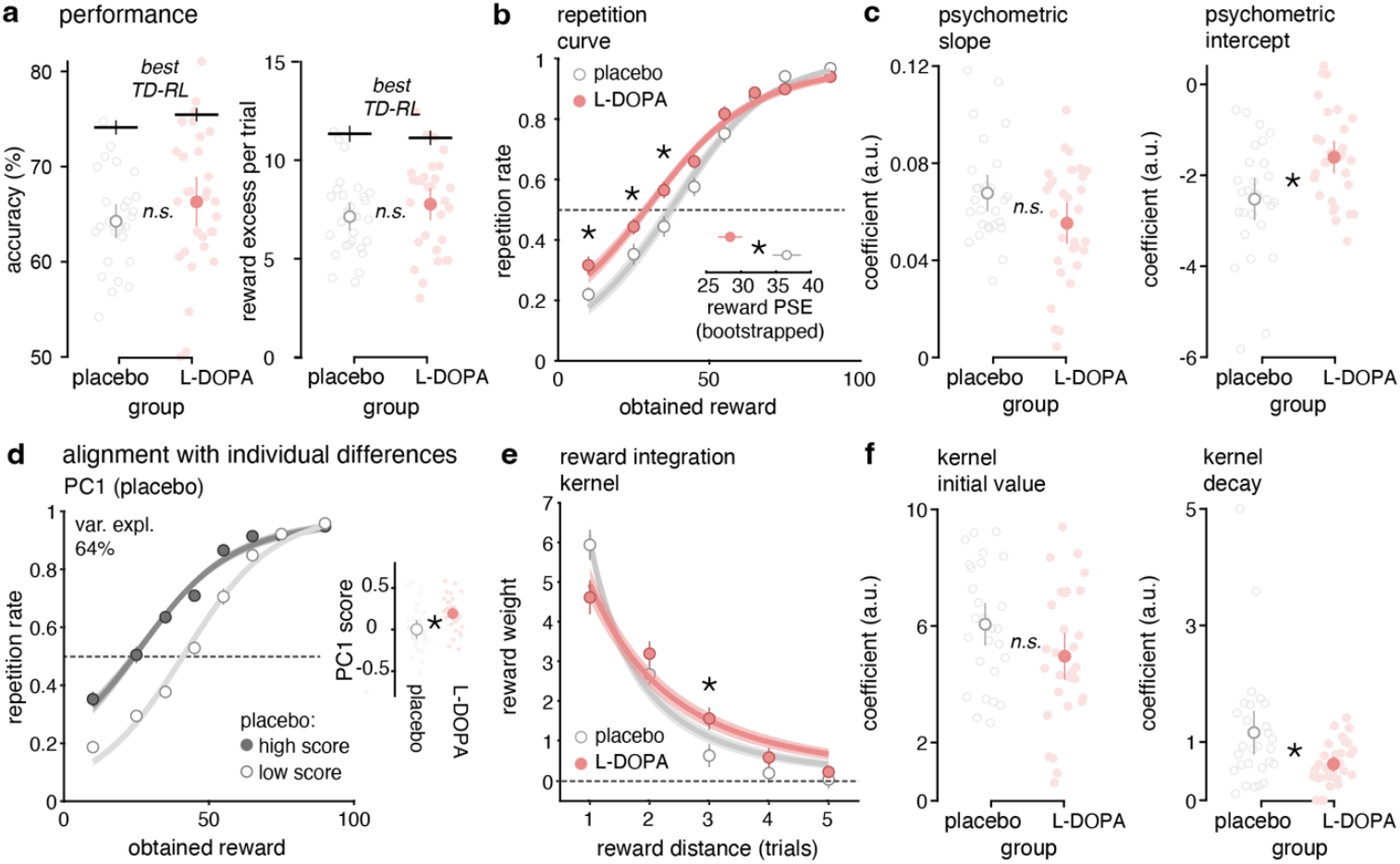
L-DOPA increases choice repetition and expands reward integration kernels. **a**. Average accuracy and excess reward per trial within each condition. Accuracy reflects whether the option selected at time *t* was the most rewarding. Excess reward represents the difference in points between the chosen and unchosen options on each trial. The dots represent individual participants’ mean performance. The two middle dots indicate the mean. Error bars represent the standard error to the mean (s.e.m.). Black horizontal bars refer to the mean and s.e.m. of accuracy and excess reward of the optimal TD-RL model. **b**. Choice repetition curves (probability of repeating the choice given the previous reward received) within each group. Points indicate the mean probability of repeating the last choice across participants. Dotted lines indicate the mean accuracy of the optimal psychometric model. Insert shows the Point of Subjective Equality (PSE-the obtained reward at which repetition rate = 0.5) extracted from the repetition curve of each participant. **c**. Slope and intercept of the best-fitting repetition curve for each participant. **d**. Repetition curve split using the median score of the first principal component (PC1) computed on the placebo group. Inserts show the score of each participant for the two groups. White and red colours correspond to the placebo and L-DOPA group respectively. Percentages indicate the variance in the probability to repeat the choice given the previous reward received explained by the first component. **e**. Reward integration kernel aligned to the participant’s current choice. Points indicate the mean weight into the logistic regression of the *n*^th^ previous element before the current choice across participants. Dotted lines indicate the mean accuracy of the optimal inverse exponential model. **f**. Kernel initial value and decay computed from the fitted inverse exponential curve for each participant. ns for not significant; \**p* <0.01.

In detailed probing of the latter effect, we found that participants under L-DOPA were more likely to repeat their previous choice following lower-than-average rewards (rank-sum test: *p* < 0.05, *z >* 2.26 for rewards below 40; all other bins: *z (57)* < 1.81, *p* > 0.05, *BF* < 1.93; **Fig. 2b**). In line with this, we found that the point of subjective equivalence (PSE—the obtained reward at which repetition rate = 0.5) differed significantly between the L-DOPA and placebo groups (rank-sum t-test: t = 122, *p* < 0.01; Fig. 2b, insert). We fitted the repetition curves with a sigmoid function to compare their slope (the parameter corresponding to the steepness of the curve) and intercept between placebo and L-DOPA groups. The intercept of the psychometric curve differed significantly between the L-DOPA and placebo groups, whereas the slope did not (rank-sum t-test: t = −1.42, *p* = 0.16 for the slope; t = 2.65, *p* < 0.01 for the intercept; **Fig. 2c & d**). This finding again support the notion that participants on L-DOPA are more likely to repeat a previous choice, rather than manifest an altered sensitivity to rewards.

To investigate whether this greater tendency to repeat choices following smaller rewards under L-DOPA reflects individual differences in the shape of the repetition curve, we conducted a Principal Component Analysis (PCA) on the placebo group’s data. For each participant, we computed the repetition curve based on obtained reward (**Fig. 2b**), resulting in a 28 participants*8 variables matrix for inclusion in the PCA. To examine how the first principal component (PC1, 64% explained variance; **Supplementary Fig. 1a**) relates to behaviour, we split the placebo group into two subgroups based on their PC1 scores and plotted the corresponding repetition curves. The resulting difference closely resembled the effect of L-DOPA, wherein participants with higher PC1 scores showed a greater tendency to repeat their choice following smaller rewards.

To confirm the above pattern, we projected the L-DOPA group onto the PCA space derived from the placebo group and compared PC1 scores between groups. This approach avoids confounding the analysis with the strong effect of L-DOPA, which would otherwise dominate the first component if both groups were included in the initial PCA. Including both groups in the PCA would have led PC1 to simply reflect the group difference, as the L-DOPA effect accounts for a large proportion of the variance in the full sample. PC1 scores were significantly higher for the L-DOPA group (rank-sum test: *p* <0.01, *z* = −2.73 for PC1; *p >* 0.05, *z* <1.01 for PC2 & PC3; **Fig. 2d** for PC1; **Supplementary Fig. 1b** for PC2 and PC3). Overall, L-DOPA lead a positive bias on repetition curve that already explains most of participant’s behaviour in the task.

### Dopamine expands reward integration kernels

Thus far, the results indicate that L-DOPA promotes a more liberal tendency to repeat choices, even following below-average outcomes. However, these analyses ignore the critical feature of integrating information across trials in a volatile learning task rather than relying on the last trial alone. To assess whether L-DOPA alters how participants integrate past outcomes over time, we estimated the decision weight of each preceding trial *t* using logistic regression. These weights reflect the extent to which each past trial influences the next choice. The decision weight associated with each previous trial in explaining the current choice decreased as a function of distance in the trial sequence from the current choice (mixed-effects ANOVA, F (4,289) = 108, *p* <0.001; Fig. 2e), in line with a learning mechanism akin to temporal-difference reinforcement learning. The placebo and L-DOPA groups showed no global difference (F (1,57) = 0.59, *p >* 0.05), meaning that L-DOPA did not influence reward weighting on average compared to placebo. However, we found a significant interaction between groups and the distance for the current choice (F (4,289) = 4.33, *p* <0.01). In effect, participants under L-DOPA weighted the most recent information less (akin to findings in Fig 2), but weighted information from more distant in time more heavily (esp. at t-3) (two-sample t-test: *p* <0.05, *z* = −2.29 for the first element; *p* <0.05, *z* = 2.31 for the third element; all other: *z(57)* <|1.25|, *p >* 0.05, *BF* <0.51; Fig. 2e). This suggests that L-DOPA engenders an extended integration of information across trials.

To test this more formally, we fitted the decay curves with an inverse exponential to compare the amplitude and decay coefficients between the placebo and L-DOPA groups. The decay constant of the inverse exponential fitted curves differed significantly between the dopamine and placebo groups, while the initial value did not reach significance (rank sum t-test: *t* = −1.81, *p* = 0.07 for the kernel initial value; *t* = 2.46, *p* = 0.014 for the kernel decay; Fig. 2f). Importantly, the kernel initial value as well as the kernel decay for the placebo and the L-DOPA groups differed significantly from 0 (one-sample t-test against 0: all *t >* 5.94, *p* <0.01 for the amplitude and decay constant, for placebo and L-DOPA groups; Fig. 2f). This is again consistent with L-DOPA rendering information integration kernels shallower, meaning they take reward information across longer time horizons into account.

### Dopamine alters two dissociable reinforcement learning mechanisms

To investigate the role of dopamine in task learning and action selection ask, we used a Temporal-Difference-based Reinforcement Learning (TD-RL) model with a Rescorla–Wagner rule to update action values^3^ (**Fig. 3a**, see **Methods**). Crucially, the updates are influenced by random noise, with the standard deviation (s.d.) set as a fraction (ζ) of the magnitude of the prediction error, as implemented in previous work^6,20,21^. Consistent with existing theories, we modelled the choice process using a stochastic ‘softmax’ action selection policy, governed by an inverse temperature, τ. Learning noise corrupts the action values that are gradually updated over time, while choice stochasticity interprets action values without modifying them, remaining independently distributed across trials, allowing us to disentangle these two sources of behavioural variability.

**Figure 3.**
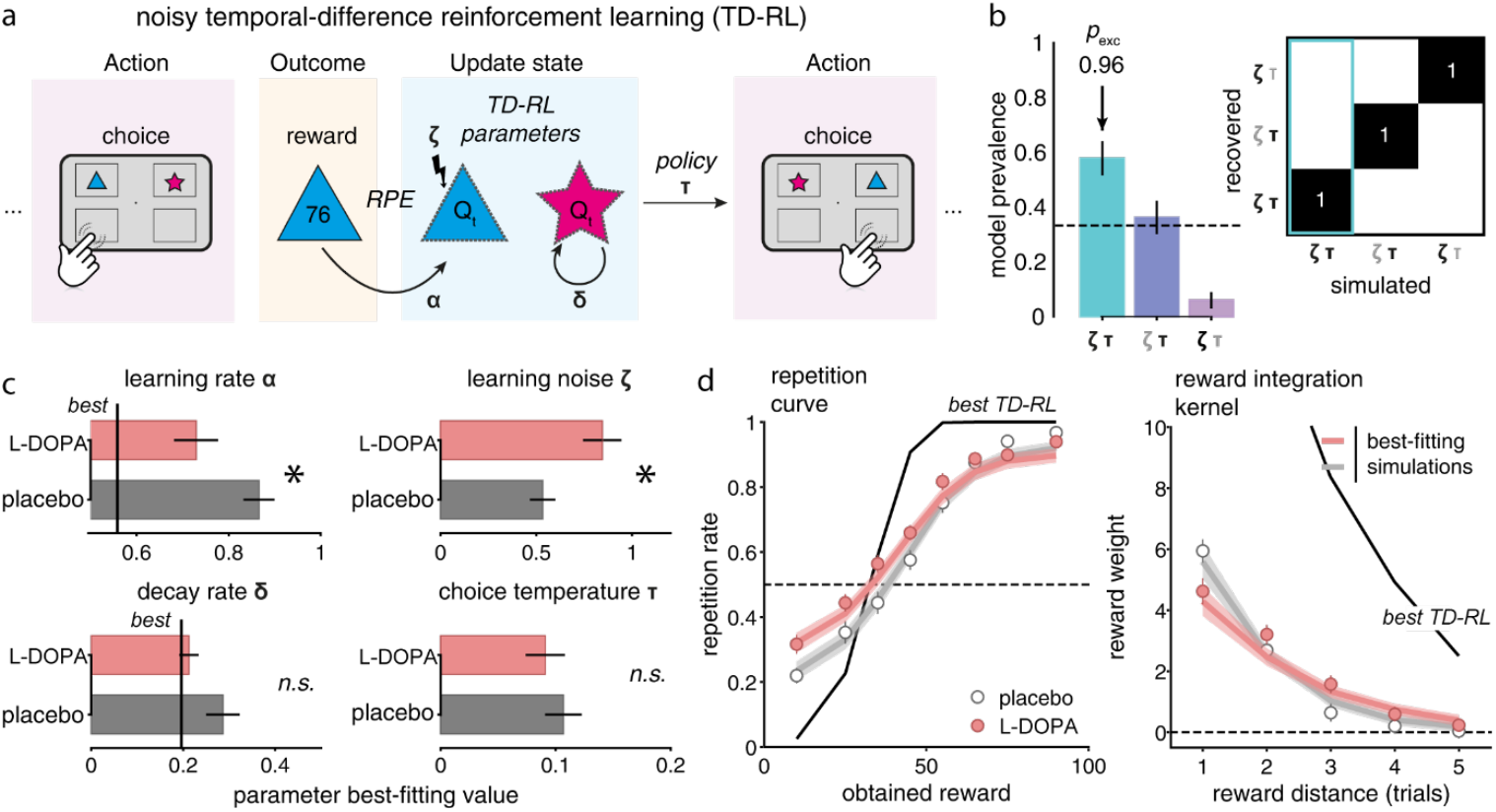
L-DOPA decreases both the rate and precision of learning in a noisy TD-RL model. **a**. Graphical representation of the noisy RL model used to fit human behaviour in the task. The Rescorla–Wagner learning rule applied to update action values is corrupted by additive random noise ζ. α represents the learning rate for the Q-value of the chosen option. δ represents a decay rate applied on the Q-value of the unchosen option. The choice process is modelled using a stochastic softmax action selection policy lead by a τ parameter. **b**. Left: Random-effects Bayesian model selection of the best-fitting strategy in participants’ data. Bars indicate the estimated model probabilities for the three candidate strategies. Model probabilities are presented as mean and s.d. of the estimated Dirichlet distribution. The dashed line corresponds to the uniform distribution. Right: Confusion matrix depicting exceedance probabilities *p*_exc_ obtained from ex ante model recovery. Greek letters in bold black indicate parameters that were free to vary in each model; those in light grey represent parameters that were fixed at zero or excluded from the models. **c**. Mean and standard error to the mean (s.e.m.) of best-fitting parameters α, ζ, δ and τ (red and grey for the L-DOPA and placebo groups, respectively) using Hierarchical Bayesian Inference (HBI). Stars indicate significant differences (*p*<.05), n.s. non-significant differences. **d**. Best fitting RL model simulations of the repetition curve and the reward integration kernel within each group. Dots indicate the mean and s.e.m. probability to repeat last choice across participants for human data. Lines indicate the mean accuracy of the optimal model.

We first evaluated whether we could distinguish between these models through model recovery (see Methods). We generated synthetic data from each model and found that we can correctly recover the ‘ground-truth’ model using Bayesian model selection (**Fig. 3b**; all exceedance probabilities > 0.83). We also ensured that the different model parameters were not correlated and that each explained distinct components of behaviour, as confirmed by parameter recovery (**Supplementary Fig. 2b**). To identify which strategy best accounted for the probability of repeating the last choice given the previous reward, and the weight of prior trials on the upcoming response, we compared three models that differed in their inclusion of choice temperature (τ) and computational learning noise (ζ), using Bayesian model selection (**Fig. 3b**; see **Methods**). In brief, one model treated both τ and ζ as free parameters; a second model implemented an argmax choice policy (i.e., choices driven exclusively by option values) while fitting ζ; and a third model treated τ as a free parameter but excluded computational learning noise (ζ fixed to zero). The noisy RL model with a softmax action policy significantly outperformed the others in explaining participants’ behaviour (exceedance probability > 0.96; **Fig. 3b**). Moreover, these two parameters were key to explaining well the behaviour of the participants (**Fig. 3b** and **Supplementary Fig. 3b & c**).

We then applied a Hierarchical Bayesian Inference (HBI) procedure^22^ to fit the selected model to participant’s data from the L-DOPA and placebo groups independently at the group level. Participants exhibited lower learning rates (α) under L-DOPA compared to placebo (L-DOPA [mean = 0. 73] vs. Placebo [mean = 0.86] group: *z* = −2.49, *p* = 0. 016; **Fig. 3c** and **Supplementary Fig. 2**). In addition, the L-DOPA group had a higher learning noise (ζ) (L-DOPA [mean = 0.84] vs. Placebo [mean = 0.53]: *z* = 2.97, *p* <0.01). No statistically significant differences were found between the L-DOPA and placebo groups for the other parameters (*p >* 0.05 for decay rate δ and choice temperature τ; **Fig. 3c**). Moreover, model simulations with a different knock-out parameter procedure (see **Supplementary Fig. 4** and **Supplementary Results**) validated the effect of L-DOPA on learning rate and computational learning noise specifically compared to the placebo group effect.

**Figure 4.**
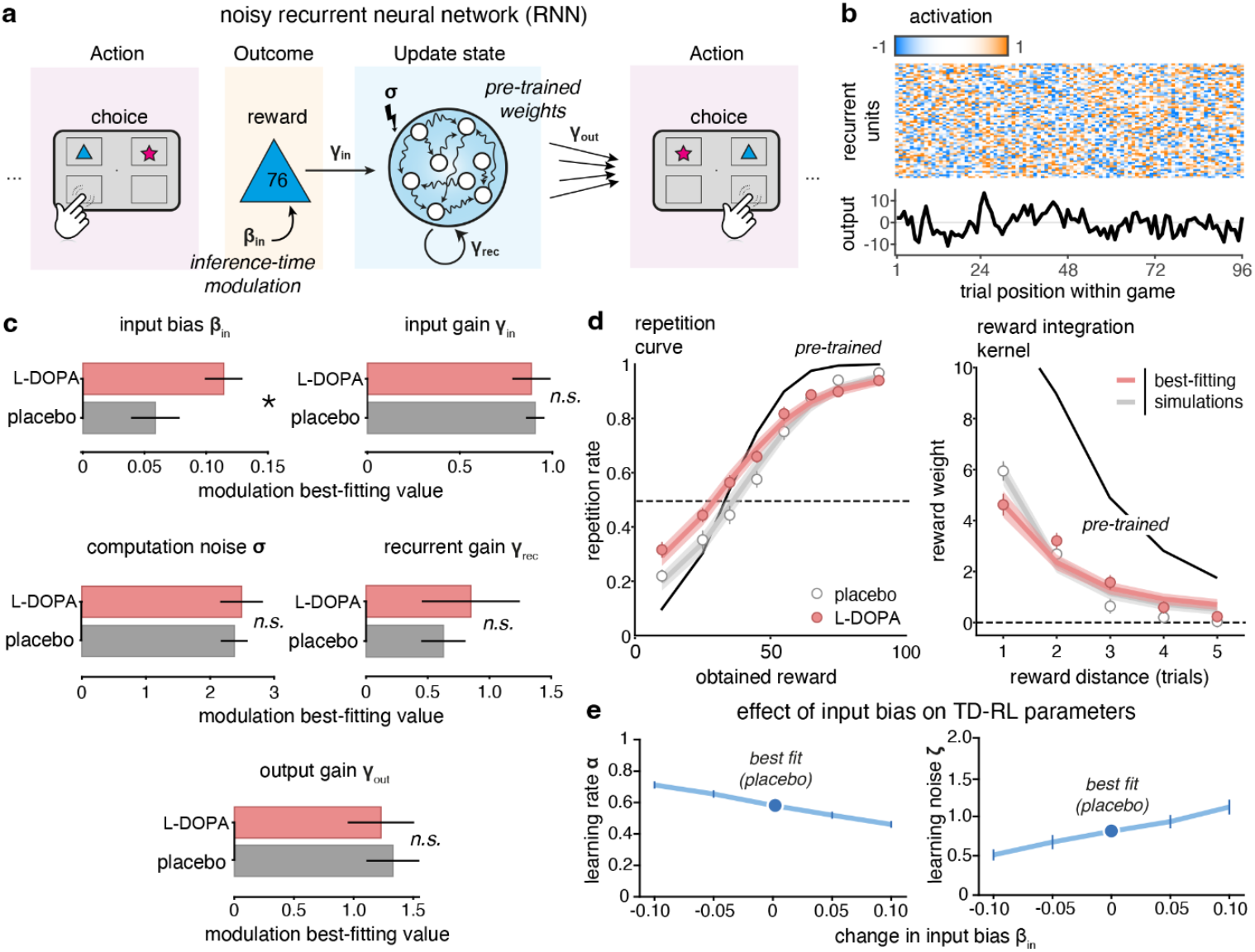
Positive input bias in noisy RNN models explains L-DOPA induced behaviour. **a**. Graphical representation of the noisy recurrent neural network model used to fit human behaviour in the task. The inputs of the optimal noisy (σ) RNN are corrupted by an additive input bias β_in_ and a multiplicator input gain γ_in_ that simulate the sustained and transient dopaminergic activity respectively. The multiplicators γ_rec_ and γ_rec_ corrupt respectively the recurrent activity and output of the RNN. **b**. Example activity of RNN recurrent units across trials during one block of the two-armed bandit task. At the bottom, the corresponding output activity across trial in favour for one option compared to another. **c**. Mean and standard error to the mean (s.e.m.) of the parameters β_in_ and γ_in_ fitted at the group level (red and grey for the L-DOPA and placebo groups respectively) using a Hierarchical Bayesian Inference (HBI) procedure. Stars represent the statistical results across groups, ns, non-significant. **d**. Best fitting RNN model simulations of repetition curve and the reward integration kernel within each group. Points indicate the mean and s.e.m. Probability to repeat last choice across participants for human data. Black lines indicate the mean accuracy of an RNN that performed the task optimally. **e**. Effect of varying RNN input bias β_in_ on the α and ζ parameters of the noisy TD-RL model, estimated by fitting the TD-RL model to RNN simulations.

It can be argued that the observed behavioural modulations—rather than being driven by a decreased learning rate (α) and increased computational noise (ζ) under L-DOPA—might instead be better explained by an asymmetric learning rate ^23,24^ or a modulation of action processing through confirmation bias ^25,26^. We fitted those two alternative models to our data to address these potential confounding explanations (see **Supplementary Fig. 5** and **Supplementary Results**). We found that an asymmetric learning rate does not account for the observed effects of L-DOPA compared to placebo. While the repetition bias parameter (ρ) improved the behavioural model fit, this more complex model still attributed the drug effects to the same mechanisms—namely, a decreased learning rate (α) and increased computational noise (ζ) under L-DOPA.

**Figure 5.**
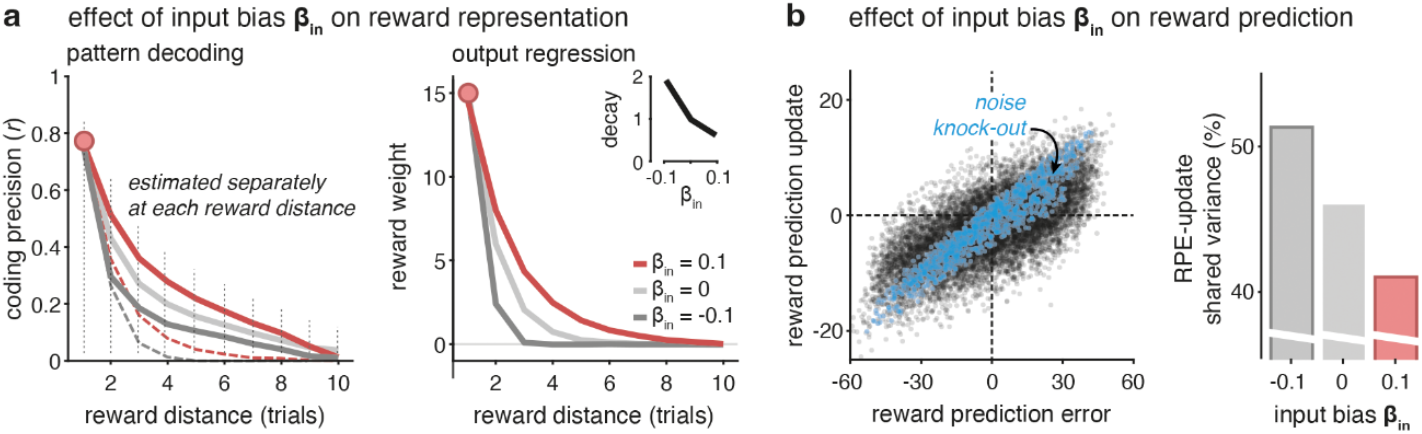
Positive input bias in noisy RNNs reproduces L-DOPA induced computational modulations. **a**. Effect of input bias β_in_ on reward representation in the RNN. Left: Reward value pattern decoding of the chosen option for the current choice and for choices up to 10 trials back (estimated separately for each trial), using the recurrent activity of the RNN at time t. Lines and shaded error bars indicate means ± sem. Dashed lines represent the same analysis but using an orthogonalized sequential decoding procedure. Right: choice-reward weight encoding (output regression) of the RNN’s recurrent activity at time t using the current choice and up to 10 preceding choices. Lines and shaded error bars indicate means ± sem. Red, light and dark grey represent results associated to an RNN with varying values of input bias β_in_ (fixed to 0.1, 0 and −0.1). **b**. Effect of input bias β_in_ on reward prediction update. Left: Reward prediction update in the best fitting RNN performing the task across trial compared to the reward prediction error. Black dots correspond to the best-fitting RNN on behavioural data; blue dots to the noise-free RNN performing the task. Right: Shared variance (r^2^ in %) between RPEs and reward prediction updates in simulated noisy RNNs across varying levels of β_in_.

### Dopamine drives a positive input bias in noisy recurrent neural networks

Next, as an implementation-level computational model, we probed the effect of L-DOPA on noisy recurrent neural networks (RNNs). We trained and tested RNNs with computational noise^19^ (**Fig. 4a**). In brief, the RNNs were trained using a reinforcement learning procedure to maximize payoff during the task (see Methods). After training, we examined whether this RNN can capture human behaviour and whether global RNN parameters capture the observed drug-induced differences. Crucially these parameters included (1) an input *bias* (β_in_) which adjusts (additive effect) the reward signal received by the RNNs; and (2) an input *gain* (γ_in_) which scale the magnitude (multiplicative effect) of the reward received by the recurrent layer (Fig. 4a). This procedure enabled an in silico test of modulation by L-DOPA of sustained (β_in_) or transient (γ_in_) dopaminergic activity, and their impact at different stages of the RNN (**Fig. 4b**). We hypothesized that an additive bias reflects sustained dopaminergic modulation, while a multiplicative gain captures transient, stimulus-related fluctuations^27,28^. It is worth noting that three additional parameters -a recurrent gain (γ_rec_), an output gain (γ_out_), and computational noise (σ) - were included in the model, but no statistically significant differences were found between the L-DOPA and placebo groups for these parameters (*p >* 0.05; **Fig. 4c** and **Supplementary Fig. 6a & b**).

**Figure 6.**
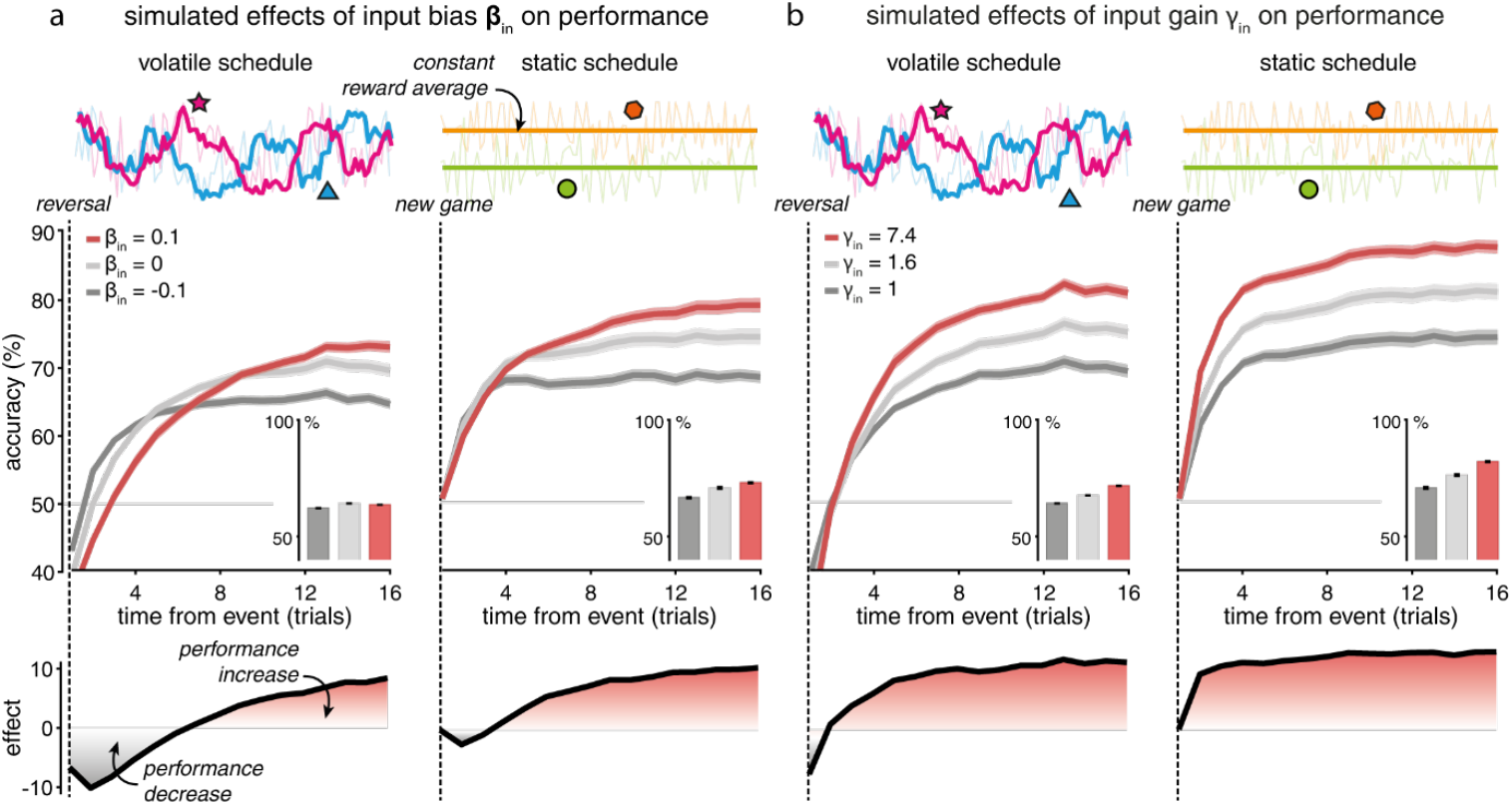
Input bias explains apparent divergence in performance between volatile and static reward schedules. Accuracy of noisy RNNs performing a two-armed bandit task with volatile (left) or static (right) reward schedules. **a**. RNNs simulated with varying input bias β_in_ values. **b**. RNNs simulated with varying input gain γ_in_ values. Inset panels depict the average accuracy of RNNs within the task. Red, light grey and dark grey curves (and bars) represent accuracy associated to noisy RNNs modulated respectively by (a) an input bias β_in_ equal to 0.1, 0 (baseline), and −0.1. or (b) an input gain γ_in_ equal to 7.4, 1.6 and 1 (baseline). Lower panels show the difference between positive and negative input modulation.

Using the same HBI procedure^22^ previously used to fit the RL model, we found an increased input bias β_in_ in the L-DOPA group (L-DOPA [mean = 0.11] vs. Placebo [mean = 0.06]: *p* = 0.011, *z(56)* = 2.63) but no change in input gain γ_in_ (L-DOPA [mean = 0.89] vs. Placebo [mean = 0.91]: *p* = 0.37, *z(56)* = −0.91, **Fig. 4c**; *p >* 0.05 for all other parameters, see **Supplementary Fig. 6a**). Moreover, model simulations and different knock-out parameter procedures (**Fig. 4d, Supplementary Fig. 7, Supplementary Fig. 8** and **Supplementary Results**) validated the input bias β_in_ as a crucial parameter to simulate in silico, using the RNNs, the effect of L-DOPA compared to placebo for this task.

### A positive input bias in RNNs explains both learning rate and noise effects

Next, to determine the association between the algorithmic and implementation level model, we tested whether a positive input bias explains L-DOPA effects on TD-RL, we simulated RNNs while varying input bias (β_in_) and fitted the simulated behaviour with the best fitting noisy TD-RL. We found that a positive input bias β_in_ on RNN computations reproduced the effect of L-DOPA on both the learning rate and precision of TD-RL (**Fig. 4e**). In other terms, applying a positive reward bias at the input of the recurrent layer implementing RL explained both L-DOPA-related effects on learning rate α and precision ζ.

While these global RNN parameters have an impact on the entire network, it was unclear how they affect activations in these networks. We first explored the impact of input modulation on reward representations in noisy RNNs, by training linear ridge regressions to predict past rewards from current RNN activations (pattern decoding approach, **Fig. 5a** and **Supplementary Fig. 9b**). and by regressing the current output activation of noisy RNNs against past rewards (output regression, **Fig. 5a** and **Supplementary Fig. 9b**). We found that a positive input bias β_in_ (**Fig. 5a**) dis not impact the strength of the representation of the last reward, but increased the representation of rewards obtained on previous trials. We also performed a sequential decoding analysis using current RNN activations and orthogonalized past rewards and found a decreased decay of the representation of past rewards up to trial t–6 (**Fig. 5a**, left, dashed lines) consistent with the output regression results (**Fig. 5a**, right). By contrast, a positive input gain γ_in_ (**Supplementary Fig. 9b**, bottom) increased the strength of the representation of the last reward and did not affect the decay of the representation of past rewards.

We then explored the impact of input modulation (input bias β_in_ and input gain γ_in_) on activations and computations of noisy RNNs during this restless bandit task. First, we tested the effect of input bias β_in_ on the output (read-out) activation of noisy RNNs and found that a positive input bias simultaneously increases the magnitude (**Supplementary Fig. 9a**, top) and variability (**Supplementary Fig. 9a**, middle) of output activation around reversals (crossings) of option values. Importantly, a negative input bias β_in_ didn’t produce the same effects, and a positive input bias increased the variability of output activation even when at a fixed magnitude of output activation (**Supplementary Fig. 9a**, bottom).

Second, we tested the impact of input modulation on the reward prediction updates assigned to each option computed by noisy RNNs. For this purpose, we trained linear ridge regressions on recurrent activations of each RNN to predict the upcoming reward of the chosen option at each trial and then computed the associated RPE corresponding to the difference between the obtained reward and the predicted reward. We also computed the associated reward prediction update, corresponding to the difference between the predicted reward for the same option at trial *t*+1 and the predicted reward at trial *t*. TD-RL hypothesizes that the reward prediction update should correspond to a fraction of the RPE, and the noisy RNNs showed the same linear positive association between RPEs and latent value updates (**Fig. 5b**, left, black dots).

To show that the strength of this association depended critically on noise in RNN computations, we computed the same variables after knocking-out noise (**Fig. 5b**, left, blue dots). As expected, the positive association between RPEs and reward prediction updates increased strongly in the absence of computation noise, showing that noisy RNNs perform noisy TD-RL computations ^6,20,21^. Modulating RNN computations using a positive input bias β_in_ of the same size as that triggered by L-DOPA decreased the strength of the positive association between RPEs and reward prediction updates, consistent with the increase in learning noise observed under L-DOPA (**Fig. 5b**, right).

Together, these analyses of RNN computations show that a positive input bias β_in_ decreases the representational decay of past rewards in RNN activations (consistent with the decreased learning rate observed under L-DOPA), while simultaneously decreasing the precision of reward prediction updates in RNN activations (consistent with the increased learning noise observed under L-DOPA).

### A positive input bias also explains dopamine effects in stationary contexts

A number of previous findings have suggested that higher dopamine levels primarily impact on learning^29^. Critically, these studies used tasks set in stable environments, whereas our task was designed under volatile conditions. To assess whether our results could align with previous findings when accounting for volatility, we simulated RNNs with increased input bias β_in_ (**Fig. 6a**) or input gain γ_in_ (**Fig. 6b**) and trained them on a related task under either volatile or stable conditions.

Examining model accuracy across conditions, we found that increasing either input gain γ_in_ or input bias β_in_ improved performance in both stable and volatile tasks. Interestingly, increasing the input bias β_in_ in the volatile condition led to higher accuracy only a few trials after a reversal. The accuracy curve crossed after four trials, resulting in no net difference in overall accuracy between different levels of input bias β_in_—mirroring the behavioural effects of L-DOPA observed in the participant’s data (**Fig.2a** for human performance and **Fig. 6a** for RNN simulations). In contrast, increasing input gain γ_in_ led to an immediate post-reversal boost in decision accuracy (**Fig. 6b**).

## Discussion

We examined the effects of dopamine administration on human reinforcement learning in volatile contexts characteristic of real-life conditions. Behaviourally, we found that L-DOPA decreases switching between volatile options following sub-average rewards. Modelling participants’ behaviour using a TD-RL model and pre-trained RNNs featuring noise in their value updates, we show a positive reward bias in RNNs reproduced the L-DOPA effect on behaviour and explained the dual effect of L-DOPA on learning rate and precision in the TD-RL model.

Previous work has reported that perturbations of dopaminergic neurotransmission – through pharmacological interventions, regulatory genes or manipulation of midbrain dopamine activity – affect the balance between repeat and switch decisions in humans and rodents^13,30–36^. We found that L-DOPA modulates this crucial component of human behaviour, consistent with previous studies of individuals with gambling disorder, a population characterized by elevated baseline dopamine release^37,38^.

Our results echo seminal work which has tied transient dopaminergic release in the ventral striatum to the computation of RPEs^10,11^ – the core signal that drives reinforcement learning. When modelling human behaviour using random noise TD-RL model in value updates^6,21^, we found that L-DOPA simultaneously decreases the rate and precision of RL. This is consistent with individual differences in these parameters across large samples of participants^18,39^. Of note, the slower learning rate observed under L-DOPA reaffirms empirical evidence linking dopamine to working memory^40–43^. Moreover, the behavioural effects of L-DOPA cannot be explained by a biased TD-RL scheme – caused by faster learning from positive than negative RPEs^24–26,44^ – as reported in humans in the presence of static reward schedules.

Dopamine has been reported as affecting action selection^45–49^ and response vigour^50,51^, by enhancing the neural representation of rewarding actions^52–54^. By accounting for noise in human TD-RL updates ^6^, we explain these apparent effects of L-DOPA on action selection by a joint reduction in the rate and precision of TD-RL. Some studies have linked mesostriatal dopamine to the updating of beliefs about hidden states in volatile environments^55–57^. Interpreting our findings through the lens of hidden-state inference, rather than purely action-selection policy, aligns with our results—specifically, the association between sustained dopaminergic activity and the learning process in our task.

TD-RL provides a widely adopted framework for studying the cognitive computations that may be altered by dopaminergic neurotransmission^10,54,58–60^. However, without precise neurophysiological recordings—available only in rare clinical populations^60^ - it is near impossible to selectively target dopaminergic networks or link specific dopaminergic activity to behaviour and cognitive computations in healthy humans. Artificial neural networks (ANNs) provide a complementary approach to TD-RL for studying neuromodulatory effects on behaviour, offering an in-silico modelling environment to test different dopaminergic modulations. The recurrence in these neural networks has been shown to be well-suited to sequential problems^61–63^, aligning well with cognitive sequence learning and TD-RL tasks. However, this optimization process relies on stochastic backpropagation algorithms that require thousands of iterations to reach asymptotic performance. Dopaminergic modulation during pharmacological intervention operates on a much faster timescale and influence already developed brain structures. To investigate its effect on learning in our protocol, we fixed the RNN weights—optimized beforehand with the same volatile reward schedule as experienced by human participants using stochastic gradient descent (SGD) — and then fitted global meta-parameters from these networks to human behaviour in the placebo and L-DOPA groups, using RNNs in a way that departs from existing approaches^64^. We found that global meta-parameters of noisy RNNs can shape learning behaviour in the task, without any retraining of the network weight structure.

We found that L-DOPA has a selective effect on RNN modulation by shifting the reward signal used as input to update action values. This positive reward bias on the RNN input produces behaviour that is best fitted using TD-RL, with a decreased rate and precision of TD-RL. By analysing the computations and latent representations used by RNNs to perform the task, we found a positive reward bias simultaneously decreases the precision of latent value updates in pre-trained networks, while triggering longerlasting representations of rewards associated with past actions in its recurrent layer. These results support the idea that L-DOPA can affect behaviour by shaping learning rather than directly influencing the action selection policy^12^. Our RNN modulation framework also offers a unique method for explaining seemingly contradictory results regarding the effects of L-DOPA. In tasks involving static reward schedules, past research has associated increased dopaminergic neurotransmission with better learning and increased perseveration^29^. Adding a positive reward bias to RNN behaviour under static conditions yields better learning with the fixed reward schedules of past research—reproducing previous behavioural results—but leads to slower (and therefore not better) learning with the volatile reward schedules used in our study.

Our results suggest that L-DOPA plays an ‘early’ role in the decision-making process, consistent with the neurophysiological effects of L-DOPA. Acute L-DOPA administration increases activation of dopamine transporters and presynaptic receptors at dopaminergic synapses. Previous studies support this mechanism, showing that acute L-DOPA administration enhances both evoked and spontaneous dopamine transmission in the substantia nigra via a presynaptic pathway^65–67^. Simulating sustained activity in RNNs also mimics increase extracellular dopamine levels, which elevate baseline dopamine concentrations below receptor activation thresholds, thereby shifting firing rates positively in response to reward delivery^27,68,69^. However, the effects of dopaminergic neurotransmission likely vary between neural circuits, with subcortical and fronto-striatal dopaminergic networks playing distinct roles in reward-guided learning^12,70^. Further studies on Parkinson’s disease– marked by basal ganglia dopamine circuit deterioration^71^ – and specific patient populations suffering from schizophrenia – associated with increased presynaptic dopamine synthesis capacity and release in striatum, alongside reduced frontal dopamine activity^72^ – could offer critical insights into the functions of these distinct systems.

Together, our findings underscore the role of increased dopaminergic neurotransmission in shaping how humans learn and behave in volatile reward environments. We show that shifting the responses to obtained rewards produce downstream effects on TD-RL and behaviour in a way that recapitulates the effects of L-DOPA across studies using different reward schedules. Using simulations of neuromodulation in artificial neural networks at ‘inference’ time, without retraining of their structure, provides a unique framework for future research in this area.

## Methods

### Participants

A total of 60 adult male participants were recruited through public advertisement in the London area. Two participants were excluded for not completing the task under the placebo condition, leaving N = 28 participants in the placebo group and 30 participants in the L-DOPA group. The recruited participants were a mean of 27,75 ± 5,9 years of age, right-handed, reported no history of neurological or psychiatric disease, and no family history of psychotic disorders. Participants reported no addiction to psychoactive drugs, nor history of psychotropic medication. Participants had normal or corrected-to-normal vision, no history of cardiac disease, and blood pressure under 140/90 mmHg. Participants provided written informed consent and received a flat £50 (plus up to £5 performance bonus) in compensation for their participation in the study. The study was approved by the University College London research ethics committee (reference: 16711/001) and all participants gave written informed consent.

### Pharmacological protocol

The study used on a double-blind, placebo-controlled, randomized protocol. Each participant performed the two-arm bandit task described below, either under Madopar CR (125 mg of L-DOPA +25 mg of Benserazide) or L-ascorbic acid (500 mg-placebo. This task was part of a neuroimaging study, and the protocol was carried out after an fMRI session. To ensure the efficacy of the dopaminergic medication throughout the 4-hour session, Madopar was used, as it contains Benserazide, a DOPA decarboxylase inhibitor that prolongs the half-life of L-DOPA. Moreover, we selected the controlled-release form of Madopar (Madopar CR) to extend the medication’s half-life. The assigned medication was dissolved in fruit squash by the attending clinician (MM or MN) and provided to the participant by the research assistant in a double-blind procedure (neither the research assistant nor the participant was aware of the medication condition). This dose oral administration was chosen in line with previous studies that used similar pharmacological protocols ^13,29,73,74^.

### Restless two-armed bandit task

We asked participants to play a restless, two-armed bandit task. In practice, participants were asked to maximize their monetary payoff by sampling repeatedly from one of two reward sources depicted by coloured shapes. Experiments were divided into two blocks of trials, each involving a new pair of coloured shapes (96 trials per block). In each trial, participants were asked to choose one of the two shapes presented to the left and right of fixation by pressing one of two buttons with their left or right index finger and then observed its associated outcome (Fig. 1b). Participants were asked to favour precision over speed, and no time limit was imposed on the latency of their responses. The payoffs that could be obtained from either shape (from 1 to 99 points) were sampled from probability distributions with means that followed a random walk-like process. More precisely, the mean payoff on trial *t* 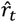 was sampled from a beta distribution with shape parameters 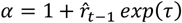 and 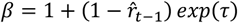 This parameterization corresponds to a mode equal to 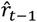 and a spread growing monotonically with τ, fixed to 3.0. The obtained payoff on trial 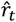 was sampled from another beta distribution with shape parameters 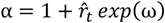 and 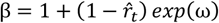 This parameterization corresponds to a mode equal to 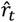 and a spread growing monotonically with ω, varied between 1.0 and 2.0 (counter-balanced across blocks). Parameter τ controls the long-term volatility of the random walk process followed by mean payoffs, whereas parameter ω controls the instantaneous uncertainty about the mean payoff 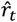 based only on the observation of the payoff *r*_t_ obtained in trial *t*. Only the obtained reward was presented—referred to as a partial outcome condition. The trajectories of mean and obtained payoffs were generated independently for each participant and each block using the procedure described above.

### Computational modelling

We derived a RL model in which the Rescorla–Wagner rule applied to update the value (expected reward) 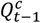 associated with the chosen action 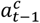 is corrupted by additive random noise ϵ_*t*_:

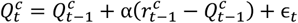

where α is the learning rate used to update action values based on the prediction error between obtained reward 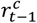 and expected reward 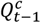 on the previous trial, and *ϵ*_t_ is drawn from a normal distribution with zero mean and s.d. σ_t_ equal to a constant fraction ζ of the magnitude of the prediction error: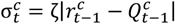 This noisy learning rule reduces to the exact (noise-free) Rescorla–Wagner rule when ζ → 0.

To test the hypothesis of the alternative asymmetric model, we dichotomized the learning rate based on positive and negative reward prediction errors (RPEs). In this model, α^+^ determines the weight given to positive RPEs, while α^-^ does the same for negative RPEs, as follows:

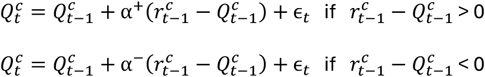

To control the decay of the value (expected reward) 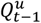 associated with the unchosen option toward the baseline value (50 points, i.e., 0.5 for values rescaled between 0 and 1) we added into the model an exponential decay factor δ:

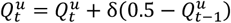

As in existing theories, we modelled the choice process using a stochastic softmax action selection policy, controlled by an inverse temperature τ:

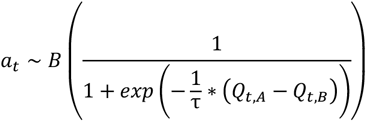

where B(.) denotes the Bernoulli distribution, and Q_t,A,_ and Q_t,B_ correspond to the values associated with actions A and B, coded as +1 for A and −1 for B. This stochastic action selection policy reduces to a purely greedy (value-maximizing) argmax policy when τ → 0. In a second alternative model, we introduced a policy repetition bias parameter (ρ), which influences the action selection process after the learning phase. Specifically, at each trial t, ρ determines the extent to which the bias *b*_t_ from the previous trial carries over to the current trial. The bias term represents a systematic tendency in action selection. At each step, this bias persists from the previous trial, weighted by ρ, such that higher values of ρ indicate stronger persistence of past biases in decision-making.

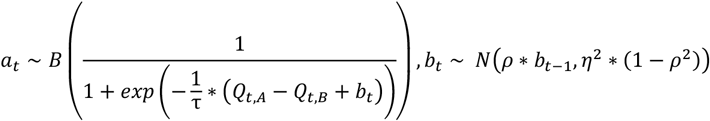

where 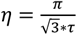 influences the stochasticity of action selection.

### Model fitting procedure for group-level parameter values

We used simulation-based methods to obtain noisy likelihoods for tested sets of parameter values, which were fed to specific fitting algorithms capable of handling such noisy likelihood functions. We fitted the candidate models to all choices using simulation-based methods, computing the response probability for each trial. We fitted the parameter values (α, δ, ζ and τ) which best explained the response probability for one option against another. We conducted model recovery analyses, detailed below, to ensure the reliability of the simulation-based fitting method outlined above. In practice, we simulated model responses (n = 1,000) to the sequences of trials presented to participants and estimated the log-likelihood of observed (participant) choice given model simulations. We combined the log-likelihood estimate with prior distributions on logtransformed parameter values to get a log-posterior estimate, whose prior distributions over parameters were defined as a truncated standard normal distributions using a range between [−10,10] (α: mean = 0 and s.d. = 1.5; δ: mean = 0 and s.d. = 1.5; ζ: mean = 0 and s.d. = 1; τ: mean = −2.3 and s.d. = 1). The unnormalized log-posterior is given as argument for the Variational Bayesian Monte Carlo (VBMC) algorithm ^75^ (version 1.0.12; https://github.com/acerbilab/vbmc) returning a variational approximation of the full posterior and a lower bound on the log-marginal likelihood. The VBMC algorithm supports stochastic estimates of the log-posterior and we provide estimates of its s.d. by bootstrapping the s.d. of the estimated log-likelihood. We finally take the posterior mean to obtain best-fitting parameter values.

The goal of this study is to compare the divergence of the fitting parameters that come from the same model between the placebo and L-DOPA group. To refine the estimation of group-level parameter values, we used the Hierarchical Bayesian Inference (HBI) algorithm^22^. In brief, we used the variational posterior of the estimated parameter to compute a prior parameter distribution for each parameter of the selected model, independently for the placebo and L-DOPA group. This distribution will then be used as parameters to build the prior functions of each model parameter for the VBMC fitting procedure. This procedure will be repeated until the divergence between two is low enough to stop it.

### Computational model validation

To ensure inclusion of learning noise (controlled by its Weber fraction ζ) and softmax choices (controlled by their temperature τ) were both necessary to fit participants’ choices, we performed random-effects Bayesian model selection based on its standard Dirichlet parameterization described in the literature^76^. In practice, as in recent works^6,18^, we compared the model including both sources of decision variability with two model variants: one without learning noise where ζ = 0, and one with a purely greedy choice policy where τ = 0. The full model outperformed the other two reduced models (exceedance of P > 0.96). Critically, we performed standard model recovery to validate our model simulation and fitting procedure^77,78^. By simulating the Rescorla-Wagner Reinforcement Learning model using individual best-fitting parameters, we established that our three candidate models were distinguishable between each other. Our fitting procedure was not biased, as demonstrated by successful model recovery: we simulated decisions from each model, then refitted all three models to this simulated behaviour for which the ground-truth model structure was known.

### Architecture of recurrent neural networks

The recurrent neural networks trained and tested here directly derived from a precedent study^20^. The artificial neural networks used correspond to standard (Elman) RNNs. Let us call *X*_t_ the input to the network, *Z*_t_ the recurrent state of the network, and *Y*_t_ the output of the network. The RNN is governed by the following equations

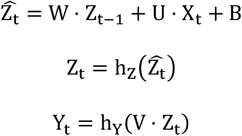

where *h*_*z*_ is the hyperbolic tangent, *h*_*y*_ is the softmax (sigmoid) function, and ⟨ · ⟩ is the matrix multiplication operator. ***W, U, B***, and ***V*** are four matrices of network parameters adjusted during training. *X_t_* is composed of the previous observed reward and a one-hot vector encoding the previous chosen action (among two possible actions). Following the first equation, the input *X_t_* and the previous recurrent activity *Z*_t-1_are integrated into an updated state 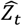 through matrix multiplications with weight matrices ***U*** and ***W*** (plus an additive bias term ***B***). This updated state 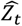 is then passed through a nonlinearity *h*_*z*_ to give the updated recurrent activity *Z*_t_. This updated recurrent activity *Z*_*t*_ then projects to action probabilities *Y*_*t*_ through matrix multiplication with output weights ***V***, followed by the softmax *h*_*y*_ operator. We used K = 64 units in the recurrent layer.

### Objective function and training procedure for noisy recurrent neural networks

The objective functions used for training the networks are derived from obtained rewards. Because the successive trials are dependent in our task, the objective function is written as follows

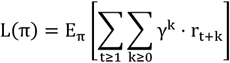

where t is the trial number, γ ∈ [0,1] is the discount factor, *r*_t+k_ is the obtained reward at trial *t* + *k* (where k is a positive integer), and π is the decision policy of the neural network. Having no prior assumptions regarding γ, we used γ = 0.5. The RNNs have a set of parameters (the matrices U, V, W, and B described in the “Neural network architecture” section) that we trained using the REINFORCE algorithm. This training procedure relies on a direct differentiation of the objective functions. The gradient of the objective function is written as follows

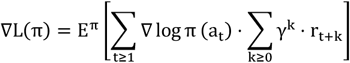

where *t* is the trial number, *a*_*t*_ is the chosen action at trial *t, r_t+k_* is the observed reward at time *t + k* (where k is a positive integer), π is the decision policy of the neural network, and γ is the discount factor. We trained 10 artificial agents using the same training procedure. The behaviour and activity patterns of the 10 trained agents were entered as repeated measures in all analyses reported below. The stochastic gradient ascent procedure was performed with the RMSProp optimizer and a learning rate of 0.0001. We set the total number of training steps to 50,000, with each step consisting of one game of 96 trials. Asymptotic performance was reached at the end of the optimization procedure.

### Introducing computation noise in recurrent neural networks

RNNs with computation noise feature random noise in the equations that govern its dynamics. We implemented computation noise in the network dynamics by updating the activity of each unit in the network in an imprecise fashion. Let *Z_t-1_* be the recurrent activity of the network at time t−1 and 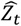 the updated state at time t before the nonlinearity *h*z The updated state of each unit in the network is corrupted with independent and identically distributed Gaussian noise. More precisely, we sampled the noisy updated state 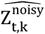 from a normal distribution of mean equal to the result of the exact (noise-free) update *Z_t,k_* and of SD σ

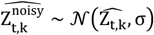

where *Z_t,k_* is the activity of unit k (k ≤ K, where K = 64 is the total number of units in the network) such that 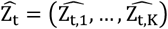 Computation noise at the unit level thus has the same structure in the two tasks. This constant-scaling noise could reflect, at least, in part, task-irrelevant input that effectively corrupts the computation of task-relevant variables. Once the noisy updated state 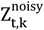 is sampled, the nonlinear activation function *h*_*z*_ is applied as in exact neural networks.

### Principal Component Analysis

PCA was performed on participants’ response reversal and response switch curves using MATLAB in-built function which centers the data and uses the singular value decomposition algorithm. Based on the first three obtained PC scores, we estimated the explained variance in reversal and switch curves through bootstrapping (n = 10,000).

### Psychometric curve fitting procedure

We fitted response repetition curves—i.e., the probability of repeating the previous response as a function of the reward received given the previous choice —by a two-parameter sigmoid function:

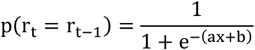

The two fitted parameters are the slope of the sigmoid curve a (indexing the sensitivity of responses to the reward received x) and the decision criterion for repeating the previous response b. As above, the bestfitting parameter values were obtained by minimizing the sum of squared errors.

### Inverse exponential curve fitting procedure

we fitted weight decisions curves by a two-parameter inverse exponential function:

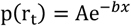

The two fitted parameters are the amplitude of the inverse exponential curve A and the decrease constant b. As above, the best fitting parameter values were obtained using the MATLAB in-built function lsqcurvefit.m. To constrain the fitting procedure and avoid aberrant fitted constant we constrained the amplitude parameter in a range of [0,50] and decrease constant between [−2,2].

### Regularized logistic regression procedure

To estimate the decision weight of each prior trial, defined as its multiplicative contribution to the upcoming choice, a multivariate parametric regularized logistic regression of choice was used. This regression relies on a linear combination of the five options chosen before the current choice:

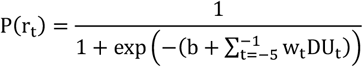

where *P*(*r*_*t*_) corresponds to the probability of selecting the current option *r*_*t*_, and b the intercept. *w_t_* represents the weight that is assigned to each of the five previous choices. These weights determine the influence each past choice has on the current decision. DU_+_ encompasses the overall utility or benefit derived from making a particular choice at that specific point in time.

### Statistical procedures

Standard paired t-tests were used to compare conditions at the group level. Data distribution (individual data points shown on the main figures) was assumed to be approximately normal and was therefore not formally tested for normality. Statistical analyses of differences between scalar metrics were performed using repeated-measures ANOVA ^79^. Bayesian Model Selection (BMS) analyses used the unbiased estimate of the marginal likelihood obtained from the model fitting procedure as the model evidence metric. This metric integrates over parameters and thus penalizes model complexity without requiring an explicit penalization term. BMS was conducted using separate fixed-effects and random-effects approaches. The fixed-effects approach assumes that all participants are relying on the same model and consists of comparing the log-marginal likelihood summed across participants for each tested model. By contrast, the random-effects approach assumes that different participants may rely on different models, and consists of estimating the Dirichlet distribution over models from which participants draw ^76^, as implemented in the SPM12 software package (Statistical Parametric Mapping).

## Acknowledgements

AZ is supported by a postdoctoral fellowship from the Fondation pour la Recherche Médicale (FRM) (SPF202209015740). VW has received funding from the European Research Council (grant agreement ERC-StG-759341). S.R.C.H. was a predoctoral fellow of the International Max Planck Research School on Computational Methods in Psychiatry and Ageing Research. The participating institutions are the Max Planck Institute for Human Development and the University College London (UCL), and funded by the Medical Research Council (MRC; grant no. MR/N013867/1). TUH was supported by a Sir Henry Dale Fellowship (211155/Z/18/Z; 211155/Z/18/B; 224051/Z/21) from Wellcome & Royal Society and a Philip Leverhulme Prize from the Leverhulme Trust (PLP-2021-040). This project has received funding from the European Research Council (ERC) under the European Union’s Horizon 2020 research and innovation program (grant agreement n° 946055). TUH is also supported by the Carl-Zeiss-Stiftung. TUH consults for limbic ltd and holds shares in the company, which is unrelated to the current project.

## Notes

### Competing Interest Statement

The authors have declared no competing interest.

## References

1. Rangel, A., Camerer, C. & Read Montague, P. A framework for studying the neurobiology of value-based decision making. (2008) doi:10.1038/nrn2357.

2. Rescorla, R. A. & Wagner, A. R. A theory of Pavlovian conditioning: The effectiveness of reinforcement and non-reinforcement. in Classical Conditioning II 64–99 (Appleton-Century-Crofts, 1972).

3. Sutton, R. S. & Barto, A. G. Reinforcement learning: an introduction. (2018).

4. Ghavamzadeh, M., Mannor, S., Pineau, J. & Tamar, A. Bayesian reinforcement learning: A survey. Found. Trends Mach. Learn. 8, 359–483 (2015).

5. Cohen, J. D., McClure, S. M. & Yu, A. J. Should I stay or should I go? How the human brain manages the trade-off between exploitation and exploration. Philos. Trans. R. Soc. B Biol. Sci. 362, 933–942 (2007).

6. Findling, C., Skvortsova, V., Dromnelle, R., Palminteri, S. & Wyart, V. Computational noise in reward-guided learning drives behavioral variability in volatile environments. Nat. Neurosci. 22, 2066–2077 (2019).

7. Wise, R. A. Dopamine, learning and motivation. Nat. Rev. Neurosci. 5, 483–494 (2004).

8. Lerner, T. N., Holloway, A. L. & Seiler, J. L. Dopamine, Updated: Reward Prediction Error and Beyond. Curr. Opin. Neurobiol. 67, 123–130 (2021).

9. Berke, J. D. What does dopamine mean? Nat. Neurosci. 21, 787–793 (2018).

10. Schultz, W., Dayan, P. & Montague, P. R. A neural substrate of prediction and reward. Science (80-.). 275, 1593–1599 (1997).

11. Schultz, W. Neuronal reward and decision signals: From theories to data. Physiol. Rev. 95, 853–951 (2015).

12. Gershman, S. J. & Uchida, N. Believing in dopamine. Nat. Rev. Neurosci. 2019 2011 20, 703–714 (2019).

13. Chakroun, K., Mathar, D., Wiehler, A. & Ganzer, F. Dopaminergic modulation of the exploration / exploitation trade-off in human decision-making. Elife 1–44 (2020).

14. Dubois, M., Habicht, J., Michely, J. & Moran, R. Human complex exploration strategies are enriched by noradrenaline-modulated heuristics. Elife 1–34 (2021).

15. Gershman, S. J. et al. Explaining dopamine through prediction errors and beyond. Nat. Neurosci. 1–11 (2024) doi:10.1038/S41593-024-01705-4.

16. Wang, Y., Lak, A., Manohar, S. G. & Bogacz, R. Dopamine encoding of novelty facilitates efficient uncertainty-driven exploration. PLoS Comput. Biol. 20, 1–27 (2024).

17. Mah, A., Golden, C. E. M. & Constantinople, C. M. Dopamine transients encode reward prediction errors independent of learning rates. Cell Rep. 43, (2024).

18. Lee, J. K., Rouault, M. & Wyart, V. Adaptive tuning of human learning and choice variability to unexpected uncertainty. Sci. Adv. 9, (2023).

19. Findling, C. & Wyart, V. Computation noise promotes cognitive resilience to adverse conditions during decision-making. bioRxiv 2020.06.10.145300 (2020) doi:10.1101/2020.06.10.145300.

20. Findling, C. & Wyart, V. Computation noise promotes zero-shot adaptation to uncertainty during decision-making in artificial neural networks. Sci. Adv. 10, 3931 (2024).

21. Findling, C. & Wyart, V. Computation noise in human learning and decision-making: origin, impact, function. Curr. Opin. Behav. Sci. 38, 124–132 (2021).

22. Piray, P., Dezfouli, A., Heskes, T., Frank, M. J. & Daw, N. D. Hierarchical Bayesian inference for concurrent model fitting and comparison for group studies. (2019) doi:10.1371/journal.pcbi.1007043.

23. Ohta, H. et al. The asymmetric learning rates of murine exploratory behavior in sparse reward environments. Neural Networks 143, 218–229 (2021).

24. Sugawara, M. & Katahira, K. Dissociation between asymmetric value updating and perseverance in human reinforcement learning. Sci. Rep. 11, 1–13 (2021).

25. Lefebvre, G., Summerfield, C. & Bogacz, R. A Normative Account of Confirmation Bias During Reinforcement Learning. Neural Comput. 34, 307–337 (2022).

26. Palminteri, S., Lefebvre, G., Kilford, E. J. & Blakemore, S. J. Confirmation bias in human reinforcement learning: Evidence from counterfactual feedback processing. PLOS Comput. Biol. 13, e1005684 (2017).

27. Liu, C., Goel, P. & Kaeser, P. S. Spatial and temporal scales of dopamine transmission. Nat. Rev. Neurosci. 22, 345–358 (2021).

28. Schultz, W. Dopamine reward prediction-error signalling: a two-component response. Nat. Rev. Neurosci. 2016 173 17, 183–195 (2016).

29. Pessiglione, M., Seymour, B., Flandin, G., Dolan, R. J. & Frith, C. D. Dopamine-dependent prediction errors underpin reward-seeking behaviour in humans. Nature 442, 1042–1045 (2006).

30. Chen, C. S., Mueller, D., Knep, E., Becket Ebitz, R. & Grissom, N. M. Dopamine and Norepinephrine Differentially Mediate the Exploration-Exploitation Tradeoff. J. Neurosci. (2024) doi:10.1523/JNEUROSCI.1194-23.2024.

31. Cinotti, F. et al. Dopamine blockade impairs the exploration-exploitation trade-off in rats. Sci. Rep. 1–14 (2019) doi:10.1038/s41598-019-43245-z.

32. Cremer, A., Kalbe, F., Müller, J. C., Wiedemann, K. & Schwabe, L. Disentangling the roles of dopamine and noradrenaline in the exploration-exploitation tradeoff during human decisionmaking. Neuropsychopharmacol. 2022 487 48, 1078–1086 (2022).

33. Kayser, A. S., Mitchell, J. M., Weinstein, D. & Frank, M. J. Dopamine, Locus of Control, and the Exploration-Exploitation Tradeoff. Neuropsychopharmacol. 2015 402 40, 454–462 (2014).

34. Strömbom, U. Effects of low doses of catecholamine receptor agonists on exploration in mice. J. Neural Transm. 37, 229–235 (1975).

35. Dongelmans, M. et al. Chronic nicotine increases midbrain dopamine neuron activity and biases individual strategies towards reduced exploration in mice. Nat. Commun. 12, 1–15 (2021).

36. Gershman, S. J. & Tzovaras, B. G. Dopaminergic genes are associated with both directed and random exploration. Neuropsychologia 120, 97–104 (2018).

37. Wiehler, A., Chakroun, K. & Peters, J. Attenuated Directed Exploration during Reinforcement Learning in Gambling Disorder. J. Neurosci. 41, 2512–2522 (2021).

38. Pettorruso, M. et al. Exploring dopaminergic transmission in gambling addiction: A systematic translational review. Neurosci. Biobehav. Rev. 119, 481–511 (2020).

39. Lee, J. K., Rouault, M. & Wyart, V. Compulsivity is linked to suboptimal choice variability but unaltered reinforcement learning under uncertainty. Nat. Ment. Heal. 2025 32 3, 229–241 (2025).

40. Luciana, M., Depue, R. A., Arbisi, P. & Leon, A. Facilitation of Working Memory in Humans by a D2 Dopamine Receptor Agonist. J. Cogn. Neurosci. 4, 58–68 (1992).

41. Westbrook, A. et al. Striatal Dopamine Can Enhance Learning, Both Fast and Slow, and Also Make it Cheaper. bioRxiv (2024) doi:10.1101/2024.02.14.580392.

42. Yoo, A. H. & Collins, A. G. E. How Working Memory and Reinforcement Learning Are Intertwined: A Cognitive, Neural, and Computational Perspective. (2021) doi:10.1162/jocn_a_01808.

43. Cools, R. & Arnsten, A. F. T. Neuromodulation of prefrontal cortex cognitive function in primates: the powerful roles of monoamines and acetylcholine. Neuropsychopharmacol. 2021 471 47, 309–328 (2021).

44. Ohta, H. et al. The asymmetric learning rates of murine exploratory behavior in sparse reward environments. Neural Networks 143, 218–229 (2021).

45. Marcelino, A. L. de A. et al. Pallidal neuromodulation of the explore/ exploit trade-off in decisionmaking. Elife 12, (2023).

46. Daw, N. D., O’Doherty, J. P., Dayan, P., Seymour, B. & Dolan, R. J. Cortical substrates for exploratory decisions in humans. Nat. 2006 4417095 441, 876–879 (2006).

47. Humphries, M. D., Khamassi, M. & Gurney, K. Dopaminergic control of the exploration-exploitation trade-off via the basal ganglia. Front. Neurosci. 6, 16922 (2012).

48. Gurney, K., Prescott, T. J. & Redgrave, P. A computational model of action selection in the basal ganglia. I. A new functional anatomy. Biol. Cybern. 84, 401–410 (2001).

49. Adams, R. A. et al. Variability in Action Selection Relates to Striatal Dopamine 2/3 Receptor Availability in Humans: A PET Neuroimaging Study Using Reinforcement Learning and Active Inference Models. Cereb. Cortex 30, 3573–3589 (2020).

50. Niv, Y. Cost, benefit, tonic, phasic: What do response rates tell us about dopamine and motivation? Ann. N. Y. Acad. Sci. 1104, 357–376 (2007).

51. Niv, Y., Daw, N. D., Joel, D. & Dayan, P. Tonic dopamine: Opportunity costs and the control of response vigor. Psychopharmacology (Berl). 191, 507–520 (2007).

52. Guitart-Masip, M. et al. Action controls dopaminergic enhancement of reward representations. Proc. Natl. Acad. Sci. U. S. A. 109, 7511–7516 (2012).

53. Guitart-Masip, M. et al. Action Dominates Valence in Anticipatory Representations in the Human Striatum and Dopaminergic Midbrain. J. Neurosci. 31, 7867–7875 (2011).

54. Coddington, L. T., Lindo, S. E. & Dudman, J. T. Mesolimbic dopamine adapts the rate of learning from action. Nature 614, (2023).

55. Starkweather, C. K., Babayan, B. M., Uchida, N. & Gershman, S. J. Dopamine reward prediction errors reflect hidden-state inference across time. Nat. Neurosci. 2017 204 20, 581–589 (2017).

56. Nour, M. M. et al. Dopaminergic basis for signaling belief updates, but not surprise, and the link to paranoia. Proc. Natl. Acad. Sci. U. S. A. 115, E10167–E10176 (2018).

57. Schwartenbeck, P., FitzGerald, T. H. B., Mathys, C., Dolan, R. & Friston, K. The Dopaminergic Midbrain Encodes the Expected Certainty about Desired Outcomes. Cereb. Cortex 25, 3434–3445 (2015).

58. Jeong, H. et al. Mesolimbic dopamine release conveys causal associations. Science (80-.). 378, (2022).

59. Kishida, K. T. et al. Subsecond dopamine fluctuations in human striatum encode superposed error signals about actual and counterfactual reward. Proc. Natl. Acad. Sci. U. S. A. 113, 200–205 (2016).

60. Batten, S. R. et al. Dopamine and serotonin in human substantia nigra track social context and value signals during economic exchange. Nat. Hum. Behav. 8, 718–728 (2024).

61. Foucault, C. & Meyniel, F. Gated recurrence enables simple and accurate sequence prediction in stochastic, changing, and structured environments. Elife 10, (2021).

62. Molano-Mazón, M. et al. Recurrent networks endowed with structural priors explain suboptimal animal behavior. Curr. Biol. 33, 622-638.e7 (2023).

63. Ji-An, L., Benna, M. K. & Mattar, M. G. Discovering cognitive strategies with tiny recurrent neural networks. Nature 1–9 (2025) doi:10.1038/s41586-025-09142-4.

64. Yang, G. R. & Molano-Mazón, M. Towards the next generation of recurrent network models for cognitive neuroscience. Curr. Opin. Neurobiol. 70, 182–192 (2021).

65. Beckstead, M. J., Grandy, D. K., Wickman, K. & Williams, J. T. Vesicular Dopamine Release Elicits an Inhibitory Postsynaptic Current in Midbrain Dopamine Neurons. Neuron 42, 939–946 (2004).

66. Gantz, S. C., Levitt, E. S., Llamosas, N., Neve, K. A. & Williams, J. T. Depression of serotonin synaptic transmission by the dopamine Precursor L-DOPA. Cell Rep. 12, 944–954 (2015).

67. Mercuri, N. B., Calabresi, P. & Bernardi, G. Responses of rat substantia nigra compacta neurones to L-DOPA. Br. J. Pharmacol. 100, 257–260 (1990).

68. Dreher, J. C. & Burnod, Y. An integrative theory of the phasic and tonic modes of dopamine modulation in the prefrontal cortex. Neural Networks 15, 583–602 (2002).

69. Grace, A. A. Phasic versus tonic dopamine release and the modulation of dopamine system responsivity: A hypothesis for the etiology of schizophrenia. Neuroscience 41, 1–24 (1991).

70. Kings, E., Ioannidis, K., Grant, J. E. & Chamberlain, S. R. A systematic review of the cognitive effects of the COMT inhibitor, tolcapone, in adult humans. CNS Spectr. 29, 166–175 (2024).

71. Poewe, W. et al. Parkinson disease. Nat. Rev. Dis. Prim. 3, 1–21 (2017).

72. McCutcheon, R. A., Abi-Dargham, A. & Howes, O. D. Schizophrenia, Dopamine and the Striatum: From Biology to Symptoms. Trends Neurosci. 42, 205–220 (2019).

73. Chakroun, K. et al. Dopamine regulates decision thresholds in human reinforcement learning in males Supplementary information. Nat. Commun. 2023 141 14, 1–14 (2023).

74. Kroemer, N. B. et al. L-DOPA reduces model-free control of behavior by attenuating the transfer of value to action. Neuroimage 186, 113–125 (2019).

75. Acerbi, L. Variational Bayesian Monte Carlo with Noisy Likelihoods. 34th Conf. Neural Inf. Process. Syst. (2020).

76. Rigoux, L., Stephan, K. E., Friston, K. J. & Daunizeau, J. Bayesian model selection for group studies - revisited. Neuroimage 84, 971–85 (2014).

77. Palminteri, S., Wyart, V. & Koechlin, E. The Importance of Falsification in Computational Cognitive Modeling. Trends Cogn. Sci. 21, 425–433 (2017).

78. Wilson, R. C. & Collins, A. G. E. Ten simple rules for the computational modeling of behavioral data. Elife 8, 1–33 (2019).

79. Caplette, L. Simple RM/Mixed ANOVA for any design - File Exchange - MATLAB Central. https://fr.mathworks.com/matlabcentral/fileexchange/64980-simple-rm-mixed-anova-for-any-design (2022).

