## Supplementary information for "Dopamine drives a positive reward bias on human reinforcement learning"

### Table of contents

| | |
|---|---|
| **Supplementary Results** | pages 2-3 |
| **Supplementary Figures 1-9** | pages 4-12 |

### Supplementary Results

**Alternative RL models.** One could argue that the observed behavioural modulations—rather than being driven by a decreased learning rate (α) and increased computational noise (ζ) under L-DOPA—might instead be better explained by an asymmetric learning rate [23,24] or a modulation of action processing through confirmation bias[25,26]. We then fitted those two alternative models to our data to address these potential confounding explanations (**Supplementary Fig. 5**).

First, we compared the best-fitting model with an asymmetric learning model by introducing a negative learning rate parameter. The observed behavioural effects—such as an increasing tendency to repeat the previous choice even when the reward was low and a reduced decay in the influence of past choices—could have been driven by asymmetrical learning of positive versus negative outcomes. However, this asymmetric model explained the behavioural participants' data less effectively than the best-fitting model (exceedance probability > 0.99 in favour of the best-fitting model; Supplementary Fig. 5a). Further analysis of the effect of L-DOPA on the asymmetric model parameters revealed a significant difference between the positive ($\alpha^+$) and negative ($\alpha^-$) learning rates (repeated-measures ANOVA, $F(1,56) = 4.77$, $p = 0.033$; **Supplementary Fig. 5a**). However, no significant interaction was found between the sign of the learning rate for the placebo versus L-DOPA groups (repeated-measures ANOVA, $F(1,56) = 2.74$, $p > 0.05$; **Supplementary Fig. 5a**). Additionally, comparing parameters between the placebo and L-DOPA groups replicated key effects observed in the best-fitting model. L-DOPA increased computational learning noise (ζ) (L-DOPA vs. Placebo: rank sum test, $z = 2.09$, $p < 0.05$; **Supplementary Fig. 5a**) and decreased the negative learning rate ($\alpha^-$) (L-DOPA vs. Placebo: rank sum test, $z = -2.25$, $p < 0.05$; **Supplementary Fig. 5a**). No significant differences were found for other parameters ($|z| < 1.5$, $p > 0.05$). These results suggest that an asymmetric learning rate does not account for the observed effects of L-DOPA compared to placebo.

We also compared the best-fitting model with a repetition bias learning model. In addition to the parameters from the best-fitting model—learning rate (α), decay rate (δ), computational learning noise (ζ), and choice temperature (τ)—this model included an additional parameter (ρ) to account for repetition bias, which is often observed in volatile reinforcement learning tasks. This hypothesis could explain the effect of L-DOPA on the likelihood of repeating a previous choice based on received rewards. The repetition bias model explained a larger fraction of participants' behavioural data than the best-fitting model (exceedance probability > 0.99 in favour of the repetition bias model; **Supplementary Fig. 5b**). Comparing parameters between the placebo and L-DOPA groups replicated key effects observed in the best-fitting model. L-DOPA increased computational learning noise (ζ) (rank sum test: $z = 2.05$, $p < 0.05$) and decreased the learning rate (α) ($z = -2.06$, $p < 0.05$). No significant differences were found for other parameters ($|z| < 1.89$, $p > 0.05$). These results suggest that while the repetition bias parameter (ρ) improves behavioural data modelling, this more complex model assigned the drug effects to the same mechanisms.

**Knock-out RL modelling.** We investigated which of the parameters implemented in the RL model drove the observed behavioural difference between the drug conditions. To do so, we applied a knock-out procedure where we scrambled the value of the parameter of interest between the two groups and then checked whether this would destroy the simulated difference in the behaviour. The learning rate α and the computational learning noise ζ both explained the behavioural difference of the participants in L-DOPA groups compared to the placebo ones. The same procedure applied for the decay rate δ or the inverse choice temperature parameter τ didn't change the group effects substantially (**Supplementary Fig. 4a & b**).

To objectively quantify these effects, we compared the difference in repetition rates between the L-DOPA and placebo groups for each knockout model with the same metric applied to participants' data (Euclidean distance of the difference between the two groups across the repetition curve). A greater divergence indicates that the corresponding parameter knockout is more critical for explaining the participants' data. We found a higher divergence for the α - ζ knock-out model compared to the other models (-75%, -29%, -4,6% respectively for $\alpha$ - $\zeta$, $\delta$, and $\tau$ knock-out models; **Supplementary Fig. 4c**). Using the same procedure, we observe similar results with the analysis of the reward integration kernel—specifically, -116%, -26%, and 19% for the $\alpha$-$\zeta$, $\delta$, and $\tau$ knock-out models, respectively (**Supplementary Fig. 4c**), again compared to the best fitting model.

**Simulations and Knock-out RNN modelling.** We investigated whether of the input bias or input gain drove the observed behavioural difference between the drug conditions. To do so, we simulated the RNNs on the task but using 3 versions of them. The first one corresponds to the RNNs modulated by $\beta_{in}$ and $\gamma_{in}$ fitted on the participants' data. The two other model variants correspond to the RNNs modulated either by $\beta_{in}$ or $\gamma_{in}$ alone. This procedure allows us to examine the effect of each parameter on the network's behaviour. We found that the repetition curve and reward integration kernel within each group were best reproduced by the RNN modulated by both fitted $\beta_{in}$ and $\gamma_{in}$, highlighting these parameters as crucial for explaining participant's behaviour. Omitting $\beta_{in}$ substantially impairs the RNN's ability to reproduce the behavioural results, whereas omitting $\gamma_{in}$ has a comparatively smaller effect (**Fig. 4d** and **Supplementary Fig. 7**). Again, to objectively quantity these effects, we compared the difference in repetition rates between the L-DOPA and placebo groups for each model and compared it to that of the best fitting model with both $\beta_{in}$ and $\gamma_{in}$. The greater the divergence, the more critical the parameter disabled is in explaining the difference between the L-DOPA and placebo groups. We found a higher divergence for the model without $\beta_{in}$ compared to the other applying this procedure to the repetition curve (-15.8 %, -67.2% and -12.2% respectively for the best fitting, fitted without $\beta_{in}$ or $\gamma_{in}$ models; **Supplementary Fig. 7c**). Using the same procedure, we observe similar results with the reward integration kernel (-36.6%, -64.1% and -32.5% respectively for the best fitting, fitted without $\beta_{in}$ or $\gamma_{in}$ models; **Supplementary Fig. 7c**).

We then investigated which parameters best captured the behavioural difference between the L-DOPA and placebo groups. To do so, we applied a knock-out procedure where we scrambled the value of the parameter of interest between the two groups then we looked at if this would destroy the difference in the behaviour between the two groups. We found that the input bias $\beta_{in}$ explained the difference in behaviour between the dopamine and placebo group but not the input gain $\gamma_{in}$ (**Supplementary Fig. 8a & b**). Finally, the projection of the vector corresponding to the repetition curve for each model on the best fitting model shows a higher divergence for the $\beta_{in}$ knock-out model compared to the other (-67%, 4.5% respectively for the $\beta_{in}$ and $\gamma_{in}$ knock-out models; **Supplementary Fig. 7c**). We observe the same results using the same procedure but using the vector corresponding to the reward integration kernel (-38%, -4.7% respectively for the $\beta_{in}$ and $\gamma_{in}$ knock-out models (**Supplementary Fig. 8c**).

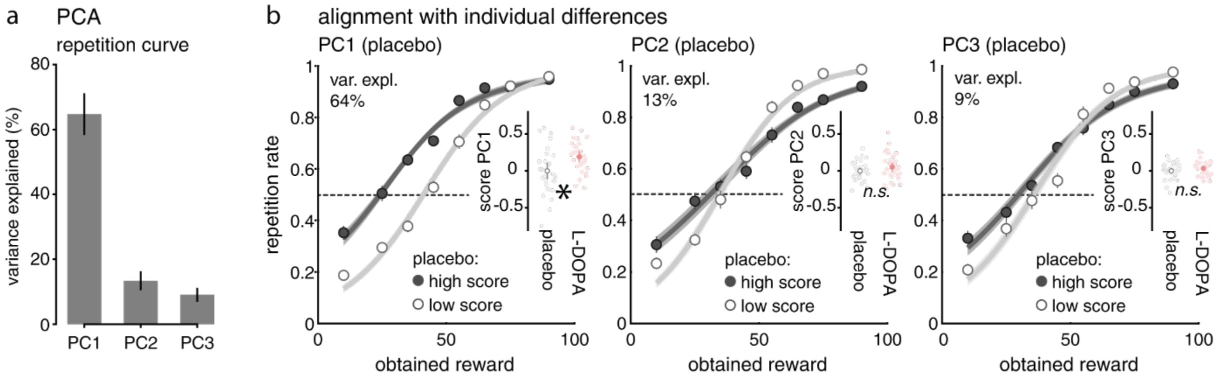


**Supplementary Figure 1. Principal Component Analysis of repetition curves. a.** Percent variance explained up to the first non-contributive PC. Black bars correspond to the mean and s.e.m. value of the percentage variance explained from the bootstrap procedure. **b.** Repetition curves for participants with high and low scores (median split) (left to right) in the first, second and third components. Inserts show the score of each participant for the two groups. White and red colours correspond to the placebo and L-DOPA group respectively. Percentages indicate the variance in the repetition rate explained by each component.

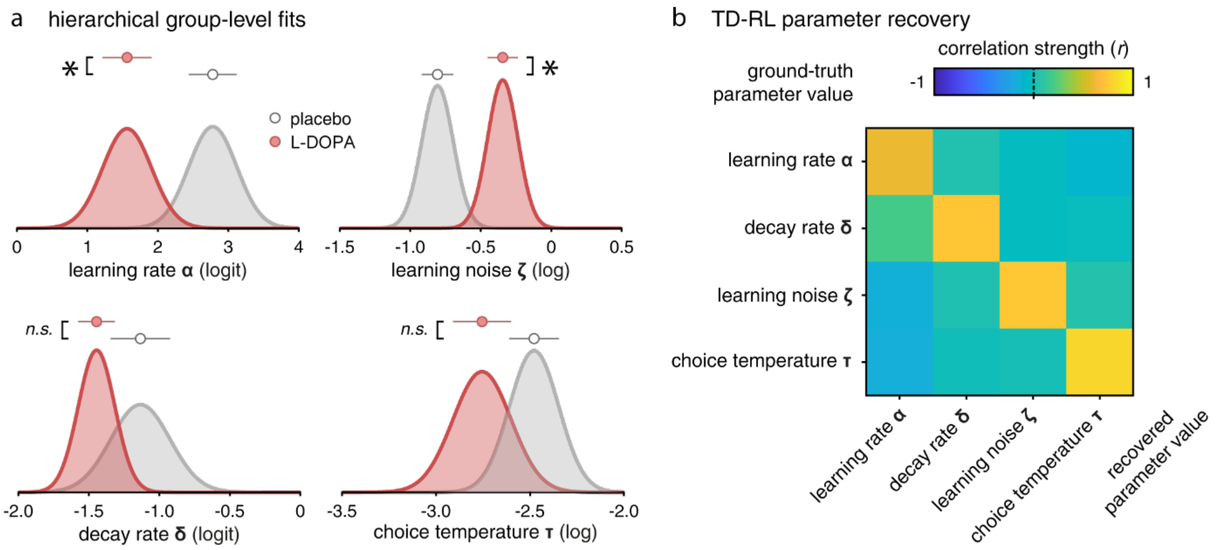


**Supplementary Figure 2. Hierarchical Bayesian Inference of group-level TD-RL parameter values. a.** Distribution of the best fitting noisy TD-RL model parameters estimated at the group level using a Hierarchical Bayesian Inference (HBI) procedure. The mean and s.e.m. for each parameter and each group are represented above the distributions. White and red colours correspond to the placebo and L-DOPA group respectively. Stars represent the statistical results across groups, ns, non-significant. **b.** Parameter recovery for the best-fitting noisy TD-RL model using a subject-level fitting procedure.

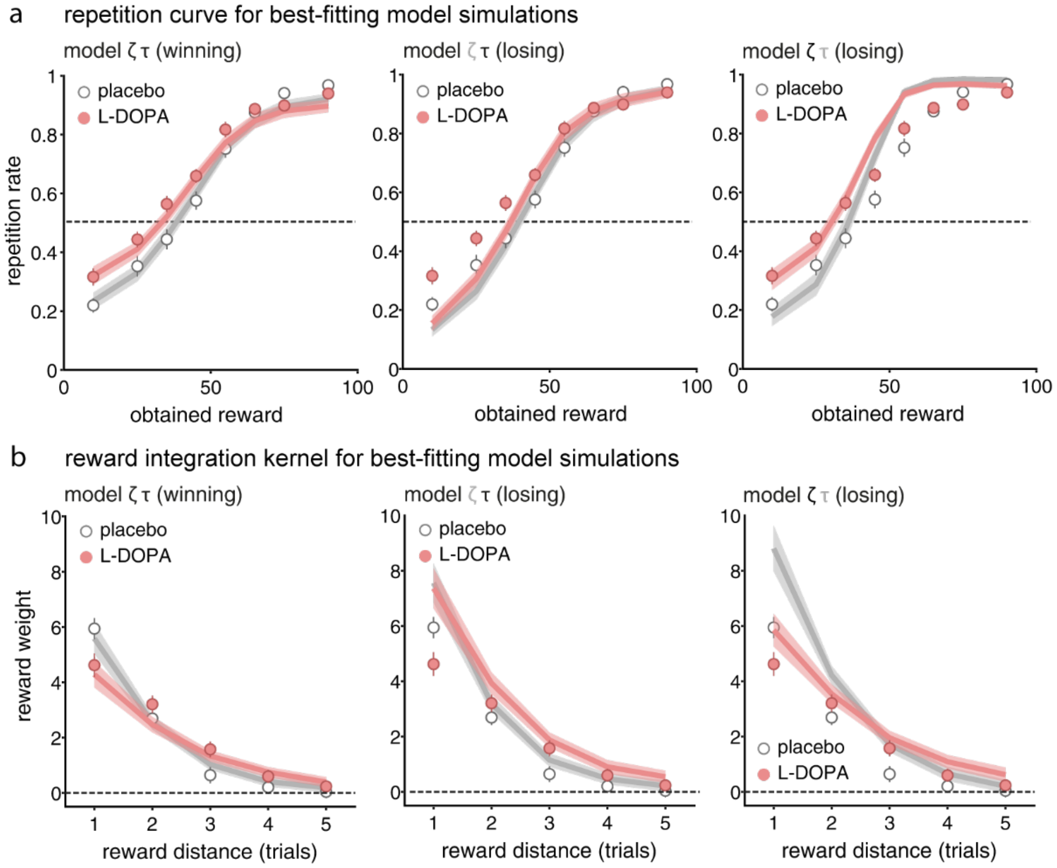


**Supplementary Figure 3. Best-fitting simulations for winning and losing TD-RL model variants.** (**a**) Simulations of the repetition curve and (**b**) the reward integration kernel within placebo and L-DOPA group, respectively (from left to right) for the best fitting model, the model without computational learning noise ζ and the model using an argmax selection policy procedure. Points indicate the mean and s.e.m. probability to repeat last choice across participants for human data. Lines indicate the mean accuracy of the optimal model.

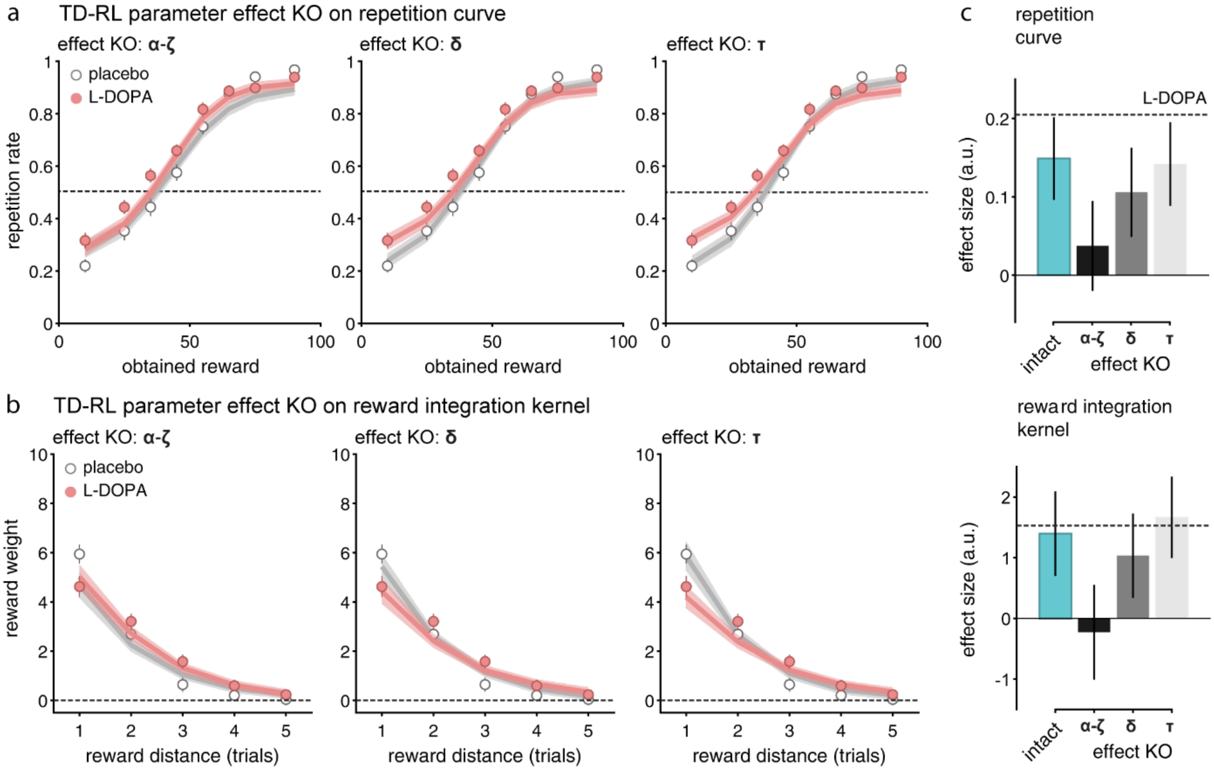


**Supplementary Figure 4. TD-RL parameter effect knock-out (KO) procedure.** Simulations of the **(a)** Repetition curves and **(b)** reward integration kernels of (left to right) α & ζ, δ and τ KO TD-RL simulations for each placebo and L-DOPA group. Points indicate the mean and s.e.m. across participants for human data. Lines indicate the mean accuracy of the optimal model. **c.** Projection of the vector representing the divergence between participants' behaviour and model simulations for the repetition curve of each model. Dashed line represents the score computed on participants' data. top: metric applied on the repetition curve. Bottom: Metric applied reward integration kernel.

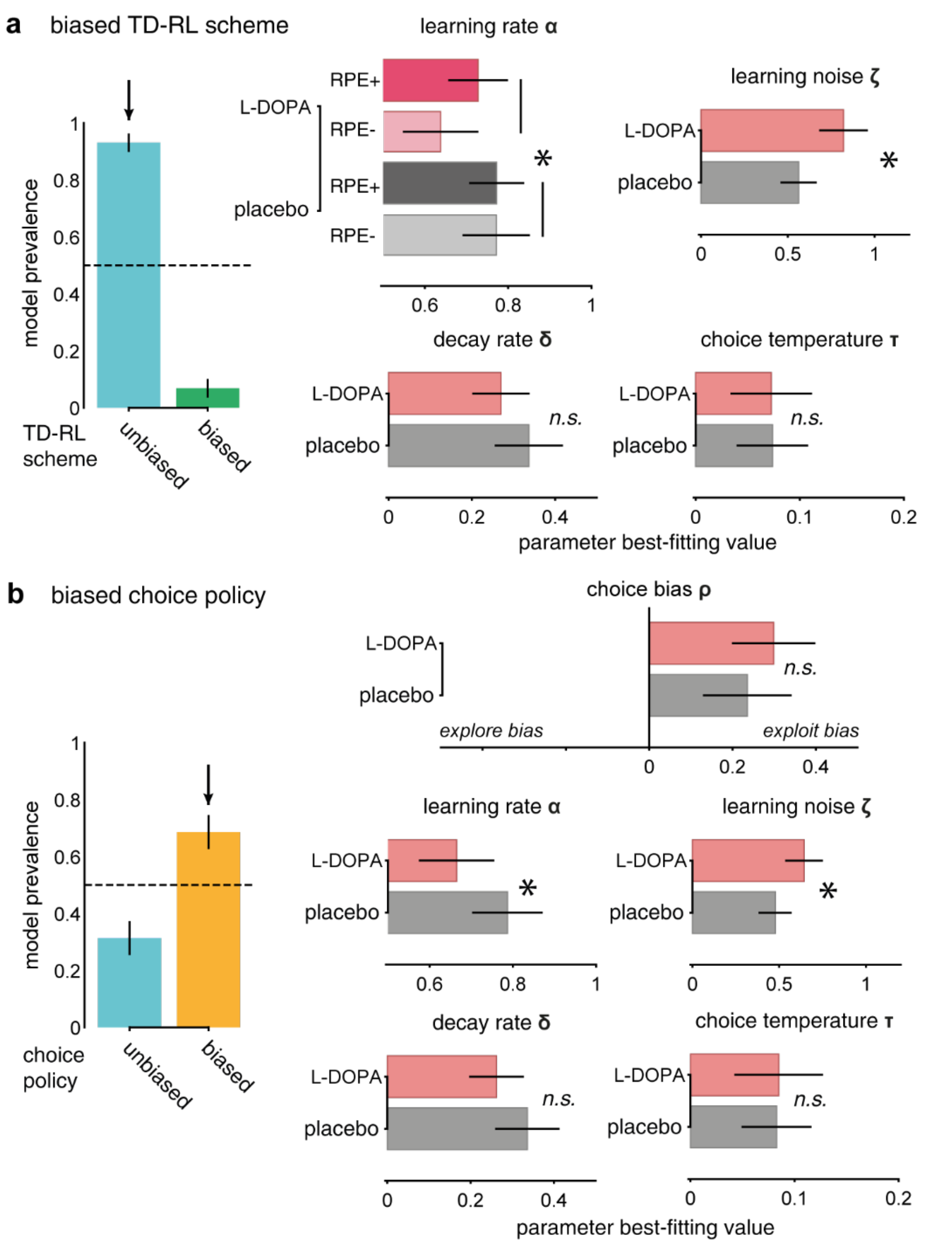


**Supplementary Figure 5. Alternative TD-RL model formulations.** Left: random-effects Bayesian model selection (BMS) of the best-fitting strategy in participants' data. Bars indicate the estimated model probabilities for the two candidate strategies. The noisy model using a softmax procedure in blue compared to the alternative hypothesis asymmetric **(a)** or policy bias **(b)** models. Model probabilities are presented as mean and s.d. of the estimated Dirichlet distribution. The dashed line corresponds to the uniform distribution**.** Right: mean and s.e.m. of **(a)** the parameters $\alpha^{+}$, $\alpha^{-}$, ζ, δ and τ and **(b)** the parameters α, ζ, ρ, δ and τ fitted by participants (red and gray for the L-DOPA and placebo groups respectively). stars represent the statistical results across groups, ns for not significant.

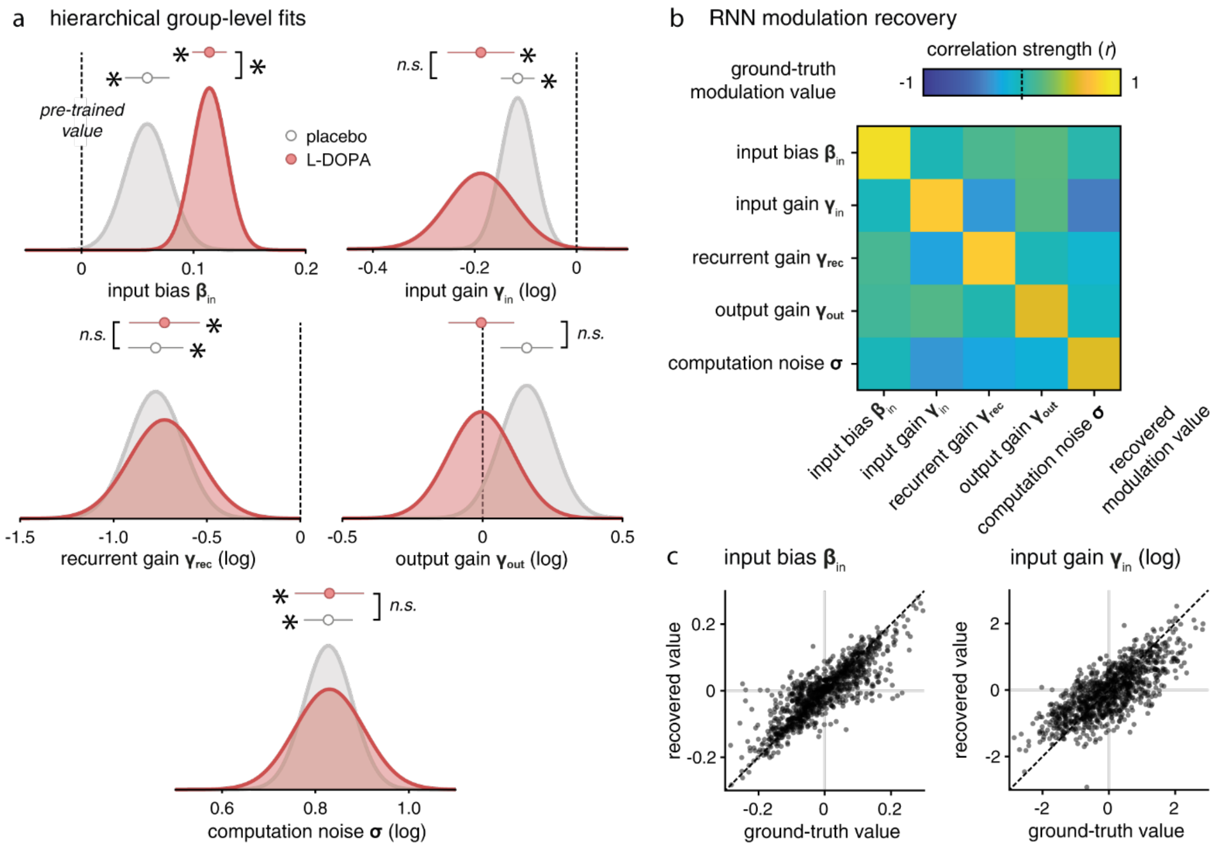


**Supplementary Figure 6. Hierarchical Bayesian Information of group-level noisy RNN modulation parameter values. a.** Distribution of the best fitting noisy RNN parameters estimated at the group level using a Hierarchical Bayesian Inference (HBI) procedure. The mean and s.e.m. for each parameter and each group are represented above the distributions. White and red colours correspond to the placebo and L-DOPA group respectively. **b.** Parameter recovery for the best-fitting noisy RNN using a subject-level fitting procedure. **c.** Correlations of ground-truth and recovered values of the input bias $\beta_{in}$ and input gain $\gamma_{in}$ parameters.

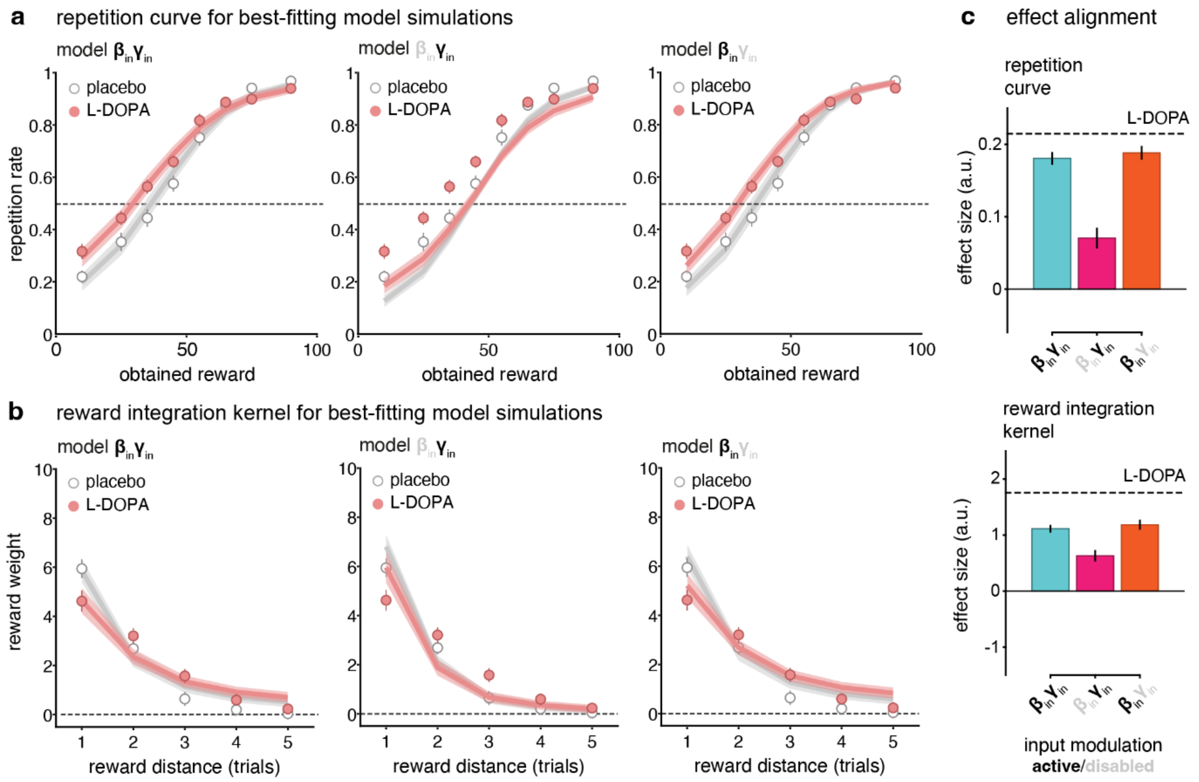


**Supplementary Figure 7. Best-fitting simulations for winning and losing noisy RNN model variants.** (**a**) Simulations of the repetition curve and (**b**) the reward integration kernel within placebo and L-DOPA group, respectively (from left to right) for the best fitting RNN model, the model without added input bias $\beta_{in}$ and the model without added input gain $\gamma_{in}$. Points indicate the mean and s.e.m. across participants for human data. Lines indicate the mean accuracy of the optimal model. **c.** Divergence between dopamine and placebo group data on the probability of repeating the last choice given the previous reward value (left) or on the reward integration kernel (right), computed for the simulated RNN models without $\beta_{in}$ or $\gamma_{in}$. The dashed line represents the score computed on participants' data.

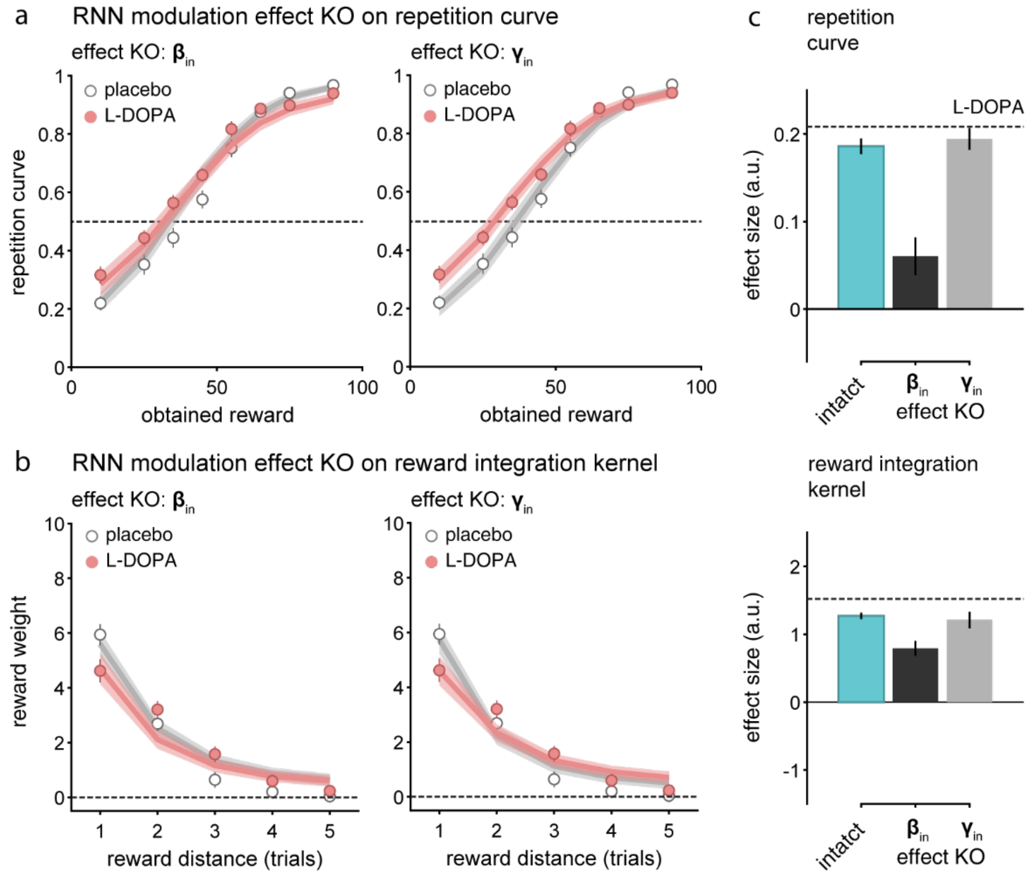


**Supplementary Figure 8. Noisy RNN modulation effect knock-out (KO) procedure.** Simulations of the **(a)** Repetition curves and **(b)** reward integration kernels of (left to right) input bias $\beta_{in}$ or input gain $\gamma_{in}$ knock-out noisy RNN simulations for each placebo and L-DOPA group. Points indicate the mean and s.e.m. across participants for human data. Lines indicate the mean accuracy of the optimal model. **c.** Projection of the vector representing the divergence between participants' behaviour and model simulations of each model. Calculated for knock-out model $\beta_{in}$ or $\gamma_{in}$. Dashed line represents the score computed on participants' data. top: metric applied on the repetition curve. Bottom: Metric applied reward integration kernel.

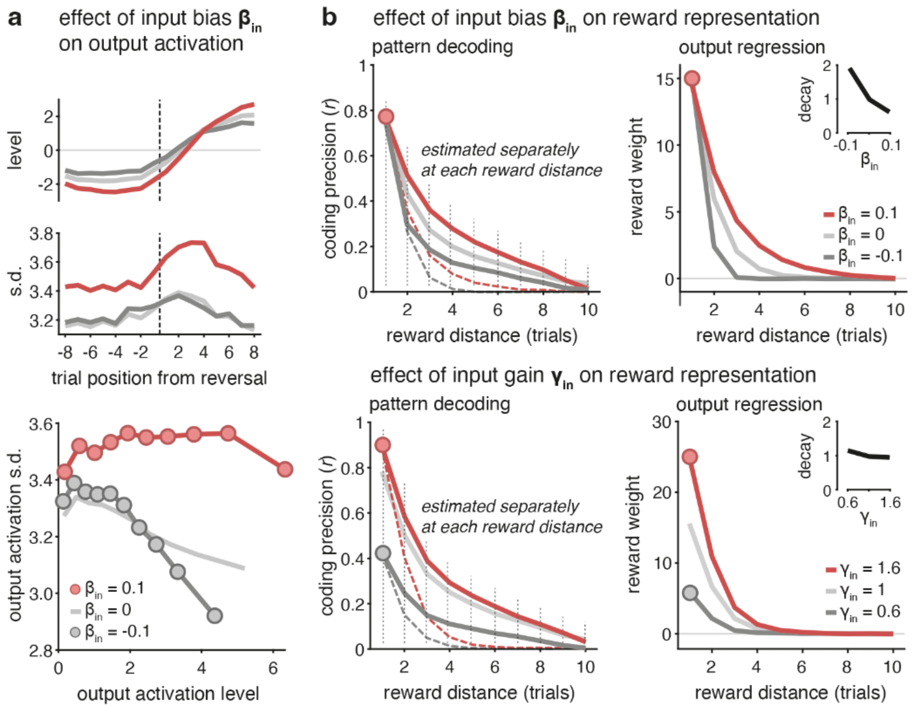


**Supplementary Figure 9. Effects of input modulation on noisy RNN activations and reward representation. a.** Effect of input bias $\beta_{in}$ on output activation of the RNNs**.** Left: mean (top) and standard deviation of the recurrent activity of the noisy RNNs performing the task given the trial position between reversal in the task. Right: Relation between mean and std of the activity of the recurrent layer of the network when playing the task. **b.** Effect of input bias $\beta_{in}$ (top) or input gain $\gamma_{in}$ (bottom) on reward representation in the RNN. Left: Reward value decoding of the chosen option for the current choice and for choices up to 10 trials back, using the recurrent activity of the RNN at time t. Lines and shaded error bars indicate means ± s.e.m. Right: choice-reward weight encoding of the RNN's recurrent activity at time t using the current choice and up to 10 preceding choices. Lines and shaded error bars indicate means ± s.e.m. Red, light and dark grey represent results associated to an RNN with value of input bias $\beta_{in}$ fixed to 0.1, 0 and -0.1 or input gain $\gamma_{in}$ fixed to 1.6, 1 and 0.6.
